# Epigenetic dysregulation of metabolic programs mediates liposarcoma cell plasticity

**DOI:** 10.1101/2025.01.20.633920

**Authors:** Erica M. Pimenta, Amanda E. Garza, Sabrina Y. Camp, Jihye Park, Samantha E. Hoffman, Laura Valderrábano, Jingxin Fu, Kevin Bi, Julie Karam, Breanna M. Titchen, Melin J. Khandekar, Erin Shannon, Yun Jee Kang, Anwesha Nag, Aaron R. Thorner, Chandrajit P. Raut, Jason L. Hornick, Priscilla Merriam, Nicole L. Solimini, Suzanne George, George D. Demetri, Eliezer M. Van Allen

## Abstract

Sarcomas are rare connective tissue cancers thought to arise from aberrant mesenchymal stem cell (MSC) differentiation. Liposarcoma (LPS) holds valuable insights into dysfunctional differentiation given its well- and dedifferentiated histologic subtypes (WDLPS, DDLPS). Despite well-established differences in histology and clinical behavior, the molecular pathways underlying each subtype are poorly understood. Here, we performed single-nucleus multiome sequencing and spatial profiling on carefully curated human LPS samples and found defects in adipocyte-specific differentiation within LPS. Loss of insulin-like growth factor 1 (IGF1) and gain of cellular programs related to early mesenchymal development and glucagon-like peptide-1 (GLP-1)-induced insulin secretion are primary features of DDLPS. IGF1 loss was associated with worse overall survival in LPS patients. Through *in vitro* stimulation of the IGF1 pathway, we identified that DDLPS cells are deficient in the adipose-specific PPARG isoform 2 (PPARG2). Defects in IGF1/PPARG2 signaling in DDLPS led to a block in differentiation that could not be fully overcome with the addition of exogenous IGF1 or the pro-adipogenic agonists to PPARG and GLP-1. However, we noted upregulation of the IGF1 receptor (IGF1R) in the setting of IGF1 deficiency, which promoted sensitivity to an IGF1R-targeted antibody-drug conjugate that may serve as a novel therapeutic strategy in LPS. In summary, lineage-specific defects in adipogenesis drive dedifferentiation in LPS and may translate into selective therapeutic targeting in this disease.

## INTRODUCTION

Soft tissue sarcomas (STS) comprise a heterogeneous collection of mesenchymal neoplasms linked to aberrant mesenchymal stem cell (MSC) differentiation. Half of all patients diagnosed with STS will develop metastatic disease, for which the median overall survival is 12-18 months(*1, 2*). The therapeutic landscape for STS overall has not evolved significantly in the past 40 years(*2*). Due to the rarity of each sarcoma subtype and the lack of representative model systems, the molecular drivers of sarcomagenesis are poorly understood. Liposarcoma (LPS) is one of the most common subtypes of STS and serves as an informative archetype to study dysregulated differentiation given its well- and dedifferentiated histologic subtypes (WDLPS and DDLPS, respectively). Both subtypes share a pathognomonic amplification of a portion of chromosome 12q inclusive of *MDM2*, but their histopathology and clinical behavior are exceptionally divergent and often co-occur within the same patient. WDLPS is histologically similar to normal adipose tissue, has a 50% local recurrence rate, and is generally thought to have no metastatic risk. In contrast, DDLPS displays a fibroblastic phenotype, has an 80% local recurrence rate, and high metastatic risk. In approximately 25% of diagnosed WDLPS cases, a DDLPS component emerges, increasing the risk of metastasis and disease-related death(*3*). Therefore, understanding the molecular mechanisms responsible for the evolution of dedifferentiation in LPS would have major clinical implications.

Previous studies have attempted to identify the drivers of the transition from WDLPS to DDLPS. Genomic analysis of paired WDLPS and DDLPS patient samples revealed no highly recurrent subtype-specific genomic events(*4*). Bulk transcriptomic studies illustrated that adipogenesis pathways are enriched in WDLPS and cell cycle programs in DDLPS(*5*), and have linked higher *MDM2* expression to an early mesenchymal phenotype(*6*); however, bulk sequencing studies are limited in their ability to resolve individual cell type specific changes. A single-cell RNA-sequencing (scRNA-seq) study investigated the cellular origins of WDLPS and DDLPS and suggested a role for transforming growth factor-beta (TGF-β) signaling within LPS subtypes(*7*). However, the scRNA-seq droplet-based strategy employed (as opposed to single-nucleus RNA-sequencing (snRNA-seq)) requires the removal of larger cells and niches of cells that are difficult to dissociate(*8*). Therefore, the larger, more differentiated WDLPS cells may have been preferentially excluded, making comparison between LPS subtypes difficult. Thus, to date, the intrinsic molecular differences between WDLPS and DDLPS tumor cells remain unknown.

In contrast to the observed aberrant differentiation in LPS, normal adipocyte differentiation begins in MSCs and is maintained through various feedback loops. Among these is an autocrine insulin-like growth factor 1 (IGF1) loop, which regulates the levels of, and signals through, the IGF1 receptor (IGF1R) to increase expression of peroxisome proliferator-activated receptor gamma (PPARG) (*9*–*13*). PPARG is a master transcription factor (TF) and regulator of adipose differentiation(*10, 14, 15*) and PPARG isoforms 1 and 2 (PPARG1 and PPARG2, respectively) are both important for adipogenesis. Low levels of PPARG1 are expressed in nearly all tissues, although under pro-adipogenic stimuli in normal preadipocytes, it induces the expression of PPARG2. PPARG2 is adipose-specific and is upregulated early in adipocyte differentiation. PPARG2 predominantly drives the expression of markers of terminal adipocyte differentiation such as *LPL, FABP4, ADIPOQ*, among others(*14, 16, 17*). While adipocytes share lineage with LPS tumor cells, whether these properties are specifically dysregulated in WDLPS or DDLPS pathogenesis are unknown.

Here, we hypothesized that distinct differentiation-specific epigenetic and transcriptomic programs of WDLPS and DDLPS underlie the biological properties driving these LPS subtypes. To address this hypothesis, we performed single-nucleus multiome-sequencing (‘multiome’; paired snRNA-seq and single-nucleus ATAC-seq (snATAC-seq)) to concurrently profile epigenetic and transcriptomic data from normal adipose, WDLPS, and DDLPS human patient tissue samples for integrated comparative analysis. We found that IGF1 chromatin accessibility, IGF1 expression, and predicted signaling through IGF1/IGF1R/PPARG were all largely absent in DDLPS. We validated these observations through orthogonal spatial profiling and projection of identified transcriptomic signatures within external LPS bulk RNA-seq cohorts. Through subsequent *in vitro* modeling, we identified that DDLPS stimulation with exogenous IGF1 did not induce PPARG2 expression. Finally, after confirming upregulation of IGF1R in the setting of dampened IGF1/PPARG2 signaling across LPS subtypes, we demonstrated IGF1R-antibody drug conjugate (ADC)-mediated selective cytotoxicity in LPS *in vitro* models. IGF1R-targeted therapies may therefore represent a novel mechanism-based selective therapeutic strategy in LPS.

## RESULTS

### Multiome analysis of LPS patient tissue samples

We performed single-nucleus multiome sequencing on WDLPS, DDLPS, and adipose patient samples (total n = 23, Figure S1A), allowing for the simultaneous characterization of both RNA sequencing and ATAC sequencing at single cell (nucleus) resolution. Pathology of the sequenced tissue was confirmed by surgical pathology reports as well as an independent pathology review of the specimens, in which all LPS samples had evidence of chromosome 12q amplification. To assess baseline biological characteristics between WDLPS and DDLPS without major confounding variables (e.g., adipose tissue depot and systemic treatment exposure), we selected patient samples that were derived from the visceral abdominopelvic or retroperitoneal regions and were predominantly treatment-naive at the time of sample collection. Patient age and sex were consistent with known LPS epidemiology (Figure S1A).

After sequencing and filtering for high quality nuclei from the snRNA-seq data, 164,229 nuclei were carried forward for downstream gene expression analysis. Cell types were annotated based on marker gene expression and inferred amplification of chromosome 12q (Figure S1B, S2), and cell types observed were consistent with those found in visceral adipose tissue^17^. snATAC-seq quality control assessments yielded 15,984 nuclei for downstream analysis. Similar cell types were recovered as seen in the snRNA-seq data, and tumor cell identity was confirmed with inferred amplifications in chromosome 12q (Figure S1C, S3). For both the snRNA-seq and snATAC-seq data, nuclei clustered according to cell type annotation and appeared to not be biased toward discernible clinical variables or confounding technical artifacts (Figures S4, S5).

### LPS subtypes have distinct transcriptomic cellular programs

Given the divergent clinical behavior between WDLPS and DDLPS tumors, we first hypothesized that distinct tumor cell transcriptional programs defined each LPS subtype. We performed differential gene expression analysis to identify differentially expressed genes (DEGs) between the WDLPS and DDLPS tumor cells. We found that IGF binding proteins, *IGF1*, and *PPARG* had significantly higher expression in WDLPS, whereas collagens (*COL1A2, COL1A1*) and *IGF1R* were highest in DDLPS (Table S1). We next evaluated enrichment of known biological pathways comprised by these subtype-specific DEGs ensuring equal patient representation and generalizability with BEANIE(*18*). Adipogenesis, insulin response, and lipid metabolism pathways were robustly enriched in WDLPS tumor cells relative to DDLPS tumor cells (Figure 1; Figure S6). Although TGF-β signaling was previously nominated as an enriched pathway in WDLPS(*7*), we observed enrichment of the TGF-β signaling pathway in only two WDLPS samples therefore this finding did not generalize across our cohort (Figure S6).

**Figure 1.**
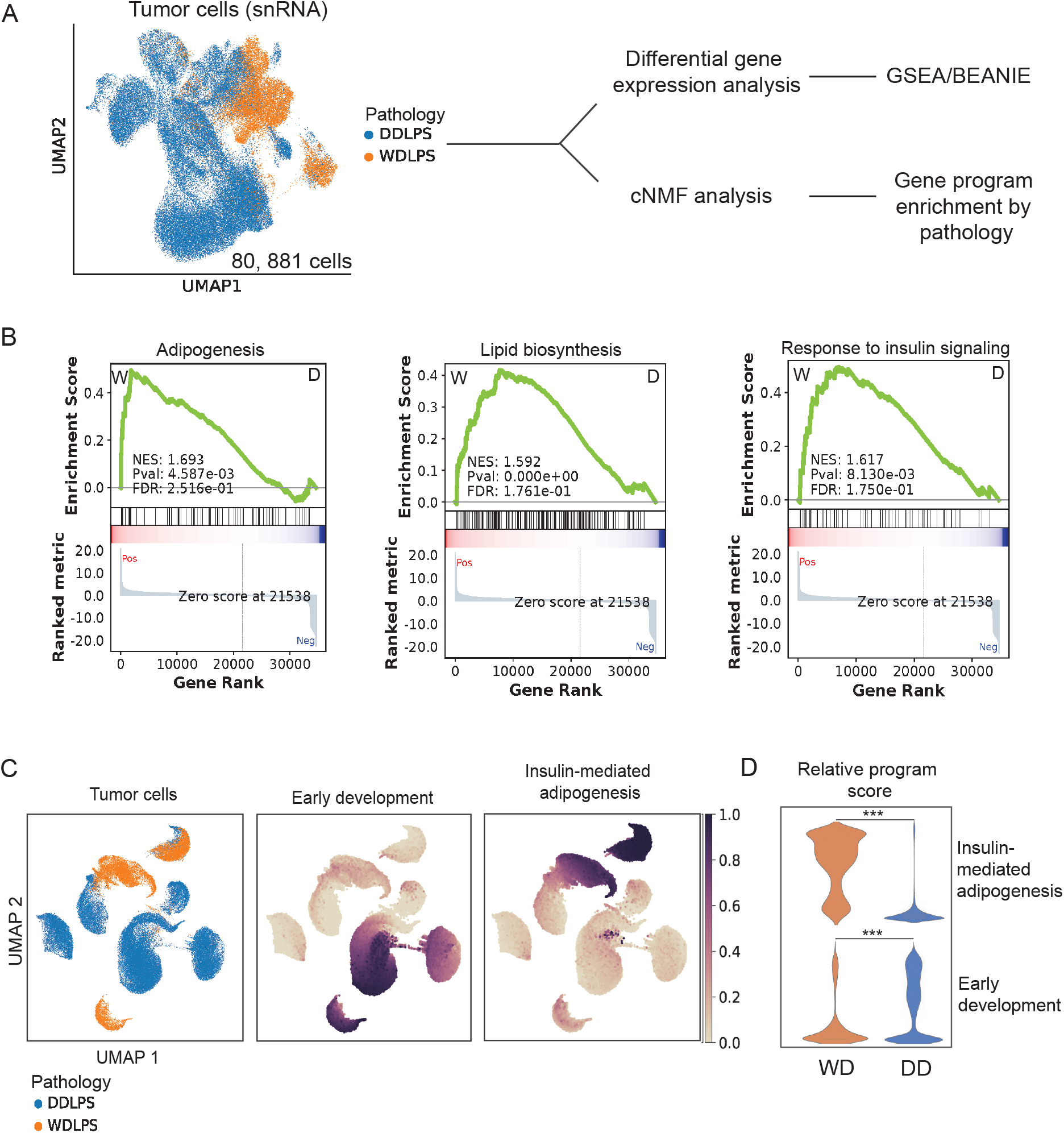
Transcriptomic analysis of WDLPS and DDLPS patient samples. (A) Schematic of the snRNA-seq analysis of LPS multiome cohort: UMAP of integrated snRNA-seq data subsetted to 80,881 tumor cells for downstream analysis, nuclei colored by LPS subtype. (B) Representative gene set enrichment plots of robustly enriched (as defined by BEANIE) gene sets in WDLPS (W) relative to DDLPS (D). (C)UMAP of unintegrated tumor cells as cNMF input colored by LPS subtype pathology (left). UMAP of tumor cells colored by relative usage of cNMF-derived cellular programs ‘Early development’ (middle) and ‘Insulin-mediated adipogenesis’ (right). (D) Violin plot of normalized usages of the cNMF programs in (C) colored by LPS subtype. *** p<0.001 by Mann-Whitney U test.

To complement these analyses, we performed consensus non-negative matrix factorization (cNMF)(*19*) to identify gene activity programs (“usage programs”) in LPS cells. Seven total usage programs were found in LPS tumor cells (Figure S7, Table S2), two of which were present across multiple patients and were significantly associated with either the WDLPS or DDLPS subtype (Figure 1C, D). The genes important to the DDLPS-associated program, termed “Early Development,” overlapped with developmental and early mesenchymal pathways involving genes such as *ROBO1, ROBO2*, and *IGF1R*. The WDLPS-associated program, termed “Insulin-mediated adipogenesis,” was composed of genes such as *PPARG* and *IGF1* and overlapped with adipocyte differentiation and insulin/receptor tyrosine kinase (RTK) signaling. Thus, both differential gene expression and cNMF analysis independently identified enrichment of genes and gene activity programs associated with insulin-mediated adipogenesis in WDLPS relative to DDLPS.

### The epigenetic landscape of LPS subtypes reveals differential IGF1 signaling regulation

The presence of histologic differences and absence of clear somatic genetic differences between WDLPS and DDLPS, paired with the identified divergence in transcriptional programming described above, led us to hypothesize that regulation of the WDLPS and DDLPS phenotype may be regulated at the epigenetic level. To probe this hypothesis, we performed differential accessibility analysis to identify differentially accessible peaks (DAPs; predicted open chromatin regions) within the snATAC-seq data in adipocytes, WDLPS, and DDLPS cells (total n = 8,873) (Figure 2A). As expected, adipocyte-specific DAPs were enriched for adipocytokine signaling, regulation of adipocyte differentiation, lipid metabolism, and insulin signaling (Figure 2B and Figure S8). DAPs in WDLPS tumor cells were enriched for IGF1 signaling, driven by peaks found within the *IGF1* and *PPARG* gene bodies. DDLPS tumor cells had differential accessibility within *RAP1B, GLP1R*, and *GNG12*, along with other genes that regulate GLP-1-mediated insulin secretion(*20*).

**Figure 2.**
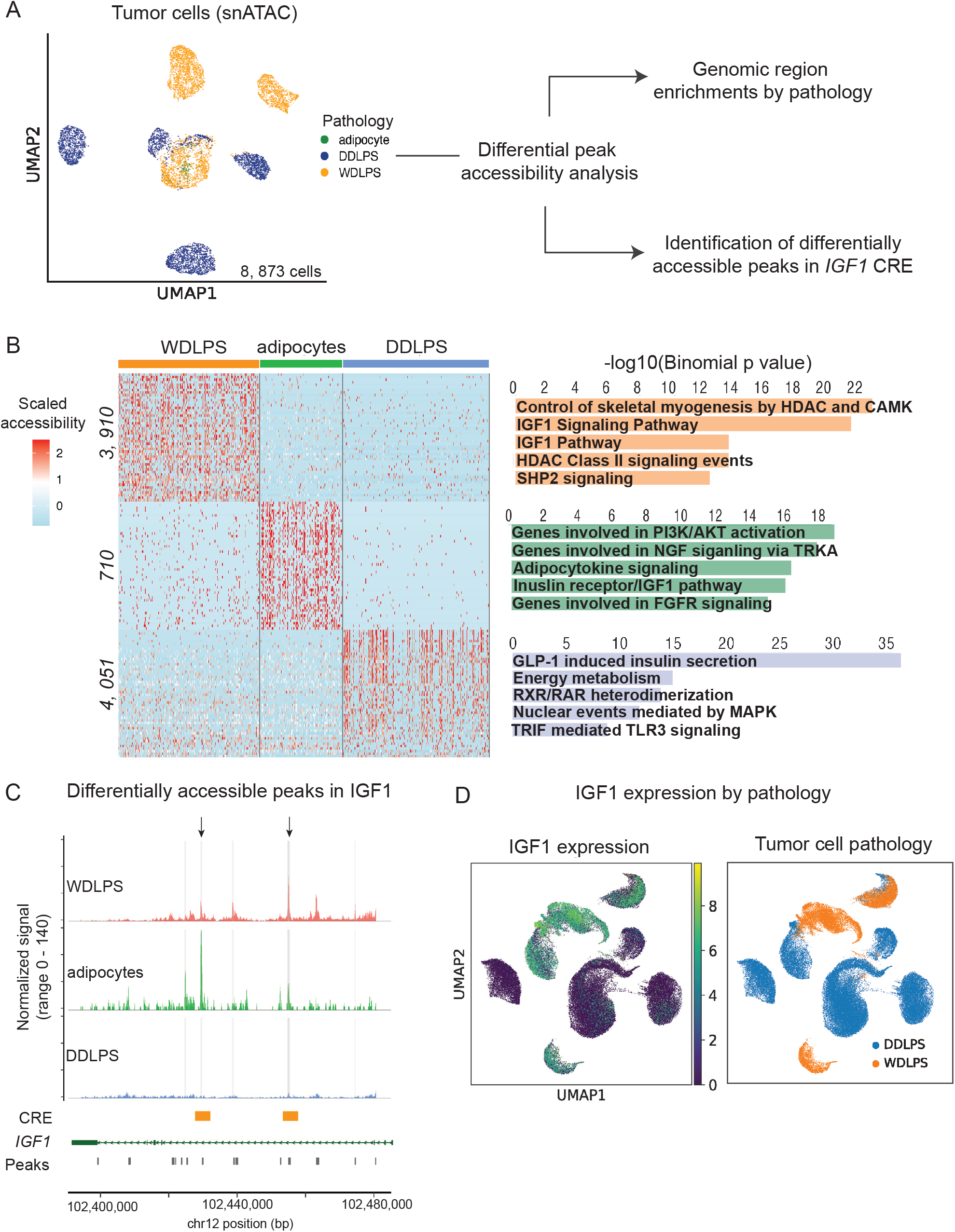
Epigenetic analysis of LPS patient tumors. (A) Schematic of epigenetic analysis of LPS snATAC-seq cohort. UMAP of subsetted 8,873 adipocytes and LPS tumor cells, colored by pathology. (B) Left, number of differentially accessible peaks per pathology and a heatmap of relative accessibility of top 250 statistically significant peaks per group. Right, top 5 significantly enriched pathways by pathology. (C) Coverage plot over IGF1 gene body by pathology. Regions highlighted in light gray are statistically significant results when using the FindAllMarkers function; regions highlighted by black arrows are significant using FindMarkers function on WDLPS and DDLPS cells only, both using logistic regression adjusted for sequencing depth. Significance defined as adjusted p value < 0.05 by logistic regression. Candidate response elements (CREs) denoted by orange bars. (D) Multiome tumor cell RNA-seq UMAP showing scaled relative *IGF1* expression (left) and tumor cell pathology (right).

Because both the transcriptomic and chromatin accessibility differences seen in WDLPS and DDLPS converged on insulin/IGF1 signaling, we next investigated whether the apparent loss of IGF1 pathway enrichment in DDLPS tumor cells was due to epigenetic silencing and subsequent expression loss of *IGF1*. We focused our analysis on only those peaks that overlapped the IGF1 gene body and calculated differential accessibility between pathologies. We observed that two peaks were significantly more accessible in WDLPS relative to DDLPS. The identified peaks were located at candidate response elements (CREs) within the *IGF1* locus (Figure 2C).

The observed *IGF1* chromatin accessibility in WDLPS and inaccessibility in DDLPS aligned with the observed *IGF1* expression levels, as *IGF1* was nearly ubiquitously expressed in WDLPS tumor cells and only expressed in approximately 10% of DDLPS tumor cells (Figure 2D). Chromatin inaccessibility over *IGF1* CREs, coupled with the loss of *IGF1* expression in most DDLPS tumor cells, suggest that *IGF1* may be an important regulator of a more well-differentiated LPS phenotype.

### Retained IGF1 expression in DDLPS is associated with better patient outcomes

As a small subset of DDLPS tumor cells retained *IGF1* expression, we subsequently hypothesized that *IGF1*-expressing LPS tumors may exhibit clinical behavior more consistent with WDLPS and therefore confer a better patient prognosis. We coupled patient outcomes data with *IGF1* expression in our snRNA-seq data and found that patients with DDLPS that retained the highest *IGF1*-expression levels remained recurrence free for more than 5 years, whereas patients with DDLPS who had lower *IGF1* expression experienced recurrence and/or disease-related death (Figure 3A). We validated this finding within the LPS TCGA bulk RNA-seq data and observed that TCGA patients with *IGF1* expression above the median had significantly longer overall survival than patients with *IGF1* expression below the median (Figure 3B). Thus, *IGF1* might orchestrate a more differentiated tumor cell transcriptional program, leading to better prognosis in patients with WDLPS and *IGF1*-expressing DDLPS tumors.

**Figure 3.**
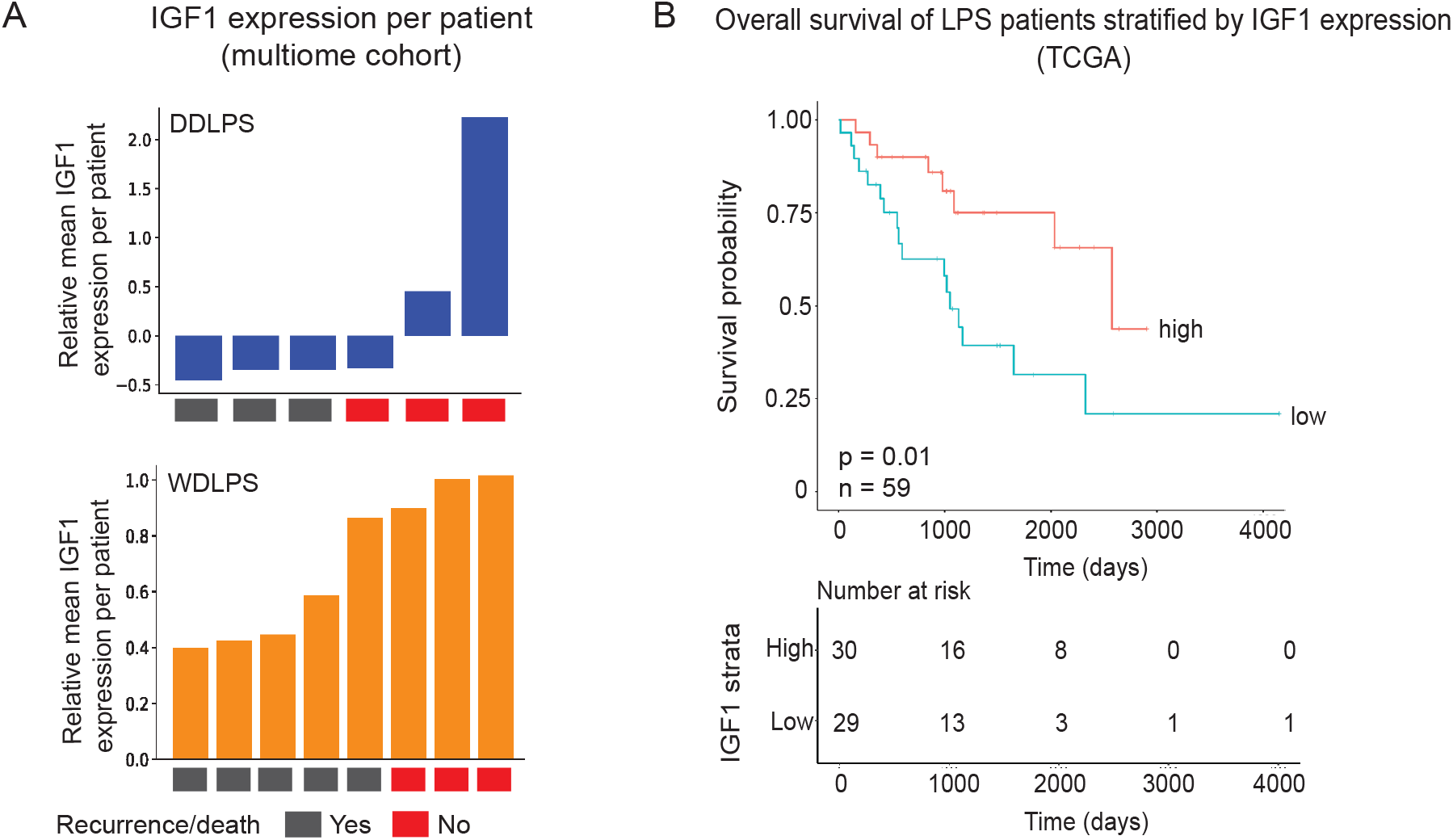
IGF1 expression and LPS patient outcomes. (A) Bar graphs of relative mean expression values of IGF1 per patient in DDLPS (blue, top) and WDLPS (orange, bottom) samples, in ascending order of expression. If a patient had two samples in the cohort, sample *IGF1* expression values were averaged. Boxes signify whether the patient has died, had recurrent LPS (gray), or has remained disease free for > 5 years (red). (B) TCGA bulk RNA-seq data comprising 58 patients with DDLPS and one patient with WDLPS was assessed for overall survival based on IGF1 expression greater than the median expression value for the cohort (‘high’) or less than (‘low’) the median. Kaplan-Meier analysis curves (top) and number at risk (bottom). Statistical significance determined by log-rank test.

### IGF1/IGF1R/PPARG signaling is absent in DDLPS

Given the above molecular findings and overall patient survival benefit of *IGF1* expression in LPS, we next sought to understand the specific mechanism by which IGF1 might affect LPS differentiation state. As a known regulator of adipogenesis, IGF1 acts in an autocrine fashion to upregulate PPARG and increase expression of downstream markers of adipocyte terminal differentiation(*11*–*13, 21*). Previous studies have also demonstrated that IGF1 negatively regulates IGF1R(*22*–*24*) (Figure 4A). Given the known importance of autocrine IGF1 signaling in maintaining adipose tissue homeostasis, we evaluated whether genes involved in IGF1 autocrine were expressed as expected in LPS. We found that normal mature adipocytes exhibited relatively high expression of *IGF1, PPARG* and markers of terminal differentiation, *FABP4* and *LPL* (Figure 4B). WDLPS tumor cells showed a similar expression pattern. In contrast, DDLPS tumor cells expressed low levels of *IGF1, FABP4*, and *LPL*, while the expression of *IGF1R* was increased more than two-fold relative to adipose tissue. These findings suggest that autocrine IGF1 signaling may be absent in DDLPS tumor cells.

**Figure 4.**
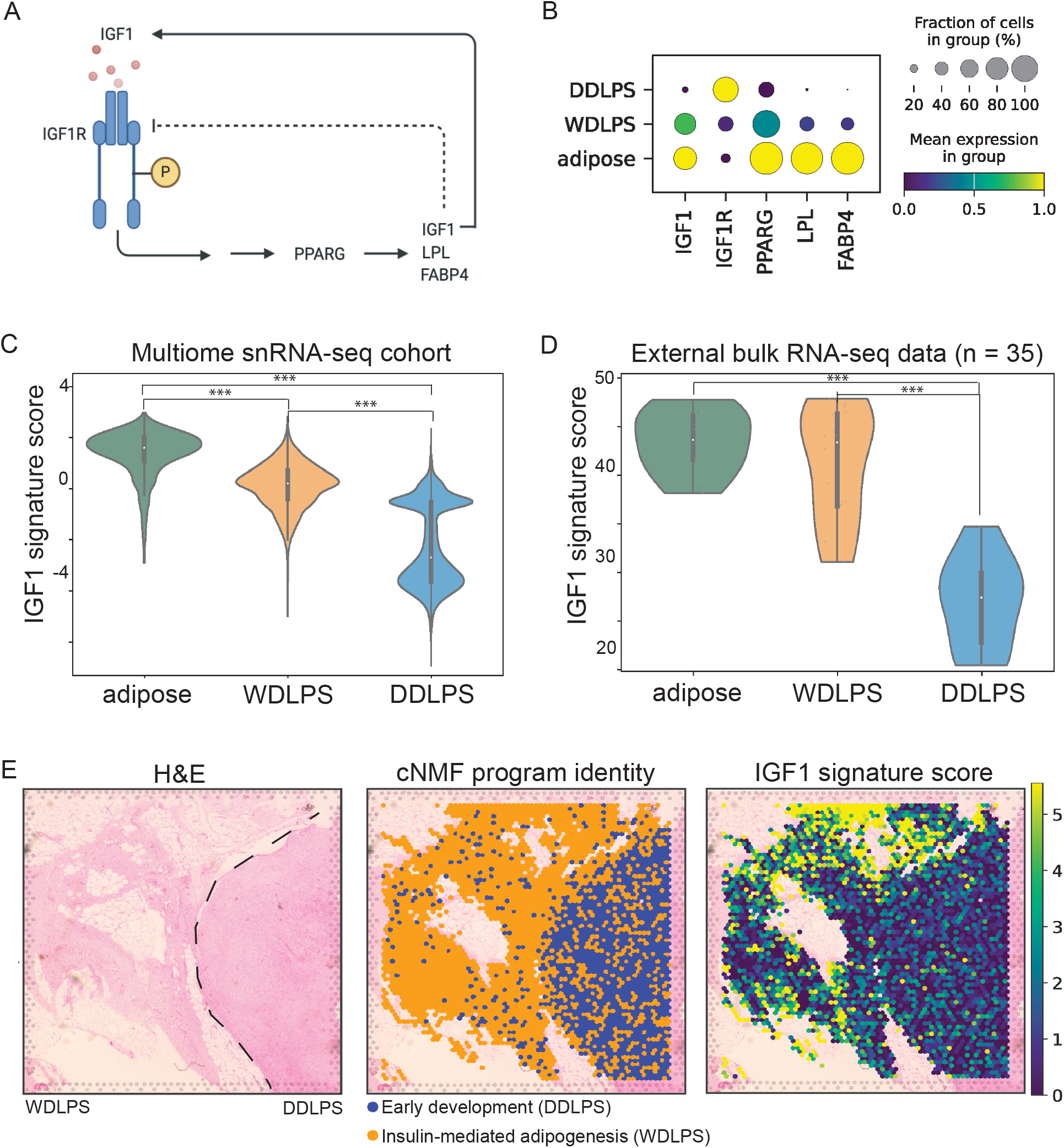
IGF1 signaling in patient LPS samples. (A) Model of adipocyte-specific autocrine IGF1 signaling. (B) Dotplot depicting scaled mean expression and percent of cells with detectable expression for each gene by LPS subtype within snRNA-seq data of multiome cohort. (C) IGF1 signaling gene signature scored within the snRNA-seq data colored by pathology. (D) IGF1 gene signature scored within external bulk RNA-seq LPS data colored by pathology. E. Spatial transcriptomic analysis of FFPE section of WDLPS/DDLPS transition zone. Left, H&E image of assessed 6.5 × 6.5 mm tissue section, LPS subtype (WDLPS, left and DDLPS, right) demarcated by dashed line. Middle, cNMF usage program with highest score assigned to each spot, colored by program name. Right, IGF1 signaling gene signature score per spot, visualized using a color scale spanning from the 10th to the 90th percentile of the calculated score, where the minimum value corresponds to the 10th percentile and the maximum to the 90th percentile *** p < 0.001 by Mann-Whitney U test.

To confirm these findings, we derived an IGF1 signaling gene signature and performed signature scoring within the snRNA-seq data (Methods). The signaling score allowed for a simplified metric to infer the relative activity of the IGF1 signaling pathway for later use across different sequencing modalities. As expected, adipocytes scored highest and DDLPS cells scored lowest for the derived IGF1 signature (Figure 4C). We subsequently evaluated the IGF1 signature in bulk RNA-seq data from an external patient cohort of normal adipose and LPS patient tissue samples(*6*) (total n = 35), demonstrating that DDLPS patient samples had a significantly lower IGF1 signaling score compared to normal adipose and WDLPS tissues (Figure 4D). Taken together, the coordinated expression and apparent loss of IGF1 signaling is a generalizable finding across DDLPS patient cohorts.

### Insulin-mediated adipogenesis and IGF1 signaling scores spatially segregate by histopathology

Given that LPS subtypes can exist as pure WDLPS tumors or as tumors that have co-occurrence of WDLPS and DDLPS components, we next hypothesized that even in the transition zone (where WDLPS and DDLPS tumor cells are spatially proximal) the derived insulin/IGF1 associated cellular programs from the multiome data would spatially segregate by histology. To probe this, we performed spatial transcriptomics at the transition zone of a mixed LPS patient tissue sample. We found that the WDLPS region of the sample was enriched for the WDLPS-associated Insulin-mediated adipogenesis cNMF usage program compared to the DDLPS region. Likewise, the WDLPS region exhibited a higher IGF1 signaling signature score relative to the DDLPS region (Figure 4E). These findings confirm that IGF1-associated tumor cell transcriptional programs appear specific to WDLPS tumor cells even in close proximity to DDLPS cells. The data imply that IGF1 may act in a tightly regulated autocrine fashion in WDLPS or that most DDLPS cells are resistant to IGF1 downstream signaling.

### IGF1-mediated in vitro adipose differentiation reveals that DDLPS cells are deficient in PPARG2

The above results suggest that IGF1 is critical in maintaining a well-differentiated adipose phenotype in LPS. We thus investigated whether exposing DDLPS cells to an IGF1-based differentiation media (IGF1-DM) was sufficient to induce adipose differentiation or whether additional defects exist in this pathway. We first selected DDLPS cell lines with transcriptional profiles similar to those of patient tumors, exhibiting low expression of *IGF1, PPARG, LPL*, and *FABP4*, but high expression of *IGF1R* (Figure S9). We cultured DDLPS-derived LPS6 tumor cells in IGF1-DM. We also cultured non-transformed human adipose-derived MSCs with IGF1-DM in parallel to serve as a positive control for the expected transcriptional response to IGF1-DM. We then compared the expression of *PPARG1, PPARG2*, and *ADIPOQ*, a marker of adipocyte terminal differentiation, at baseline and after differentiation induction. At baseline, DDLPS cells exhibited lower levels of *PPARG1* and *PPARG2* relative to the undifferentiated MSCs (Figure S10). After inducing differentiation, we observed a lack of response to IGF1-DM in DDLPS cells relative to MSCs, as evidenced by the difference in induction of *ADIPOQ*. In MSCs, *ADIPOQ* expression increased by more than 10,000-fold relative to baseline, largely due to the increase in *PPARG2* upstream. In DDLPS cells, neither *PPARG2* or *ADIPOQ* significantly increased expression in response to IGF1-DM (Figure 5A). These data are consistent with previous work illustrating that PPARG2 is the major PPARG isoform involved in the terminal differentiation(*25*–*27*) of normal MSCs to adipocytes and suggest that PPARG2 is both deficient and not inducible in DDLPS cells.

**Figure 5.**
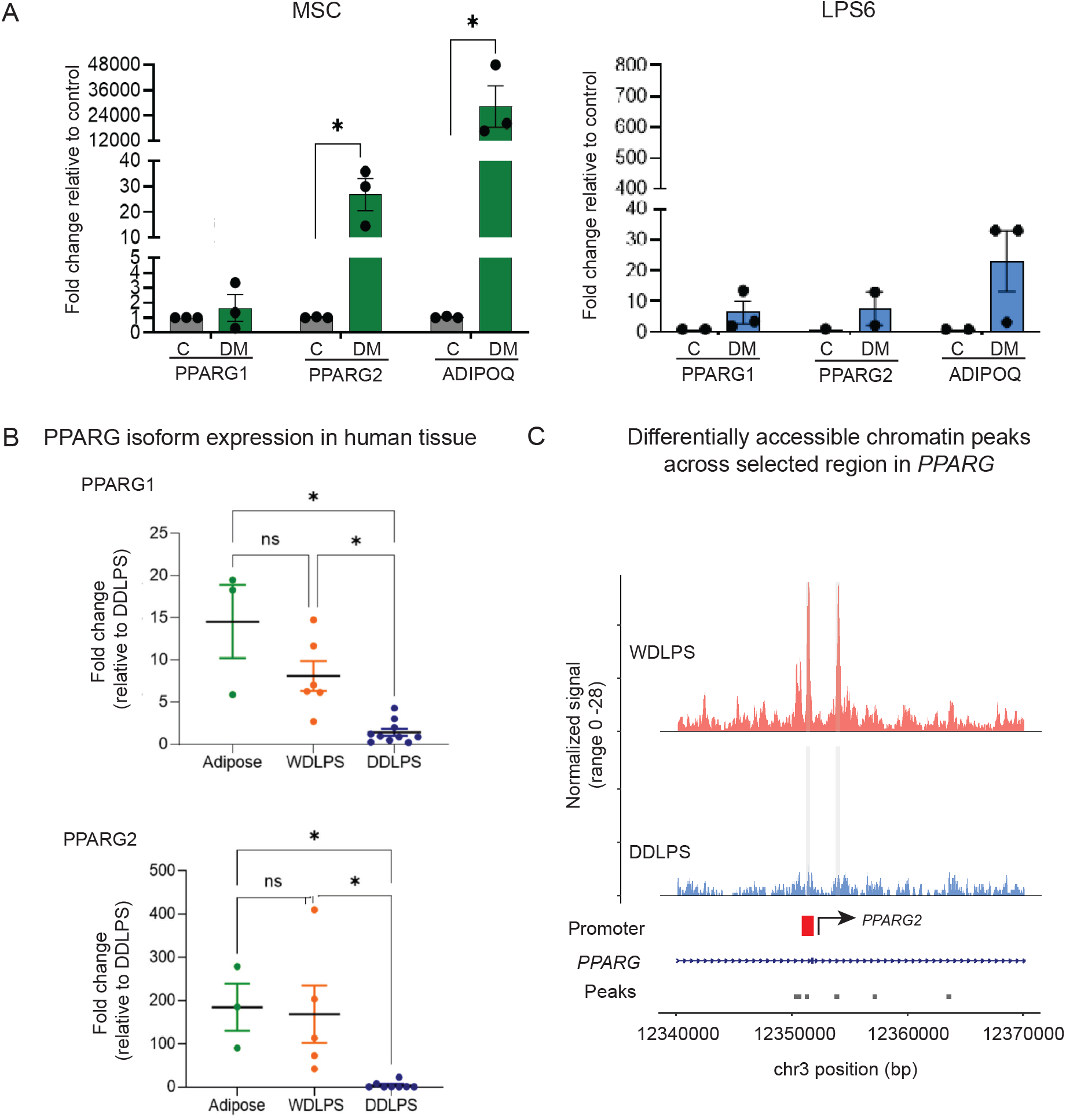
PPARG isoform expression in distinct differentiation states *in vitro* and in LPS patient tissue samples. (A) Expression of *PPARG1, PPARG2*, and *ADIPOQ* in MSCs before (control, gray) and after exposure to IGF1+DM (DM, green). On the right, expression of *PPARG1, PPARG2*, and *ADIPOQ* in LPS6 cells before (C, gray) and after IGF1+DM exposure (DM, blue). Gene expression levels are fold change delta-delta CT, normalized to beta-actin and relative to control for each cell line. Statistical significance determined by Student’s t-test. * p < 0.05. (B) Expression of *PPARG1* (top) and *PPARG2* (bottom) in human adipose (green, n = 3), WDLPS (orange, n = 6), and DDLPS (blue, n = 10) tissue samples (total n = 19; 12 samples are derived from adjacent tissue of tumors sequenced in multiome cohort, augmented by 7 additional patient samples). *PPARG2* transcript was not detected in one WDLPS and two DDLPS samples. Statistical significance defined as p < 0.05 by Kruskal Wallis. (C) Coverage plot over selected region of *PPARG* gene body containing differentially accessible peaks (light gray) in the snATAC-seq multiome data, colored by LPS subtype. Peaks deemed significantly differentially accessible between WDLPS and DDLPS cells by logistic regression, adjusted for sequencing depth. Significance defined as adjusted p value < 0.05. Promoter-like signature region (‘Promoter’) denoted by the red bar. *PPARG2* transcription start site indicated by arrow.

To ensure that the observed deficit of PPARG2 in DDLPS cell lines was relevant to human LPS tissue, we performed QPCR analysis to assess the expression of PPARG isoforms in human normal adipose, WDLPS, and DDLPS tissue samples (n = 19; Methods). We observed that the expression of *PPARG1* was about eight-fold higher in adipose and WDLPS tissue relative to DDLPS. The expression of *PPARG2* was nearly 200-fold higher in adipose and WDLPS tissue relative to DDLPS (Figure 5B). These data confirm that loss of *PPARG2* expression is a feature of human DDLPS.

We next hypothesized that loss of PPARG2 in DDLPS may be due to epigenetic silencing over the *PPARG* isoform 2 transcription start site (TSS). Analysis of the snATAC-seq data illustrated that there are two DAPs within the *PPARG* gene body between WDLPS and DDLPS (Figure S11 and Figure 5C). The observed DAPs indicate that WDLPS cells had accessible chromatin over a promoter-like sequence adjacent to the TSS of *PPARG2*. This same region was inaccessible in DDLPS cells. These data suggest that chromatin compaction over a promoter region proximal to the *PPARG2* TSS may contribute to the lack of *PPARG2* expression in DDLPS and a diminished capacity to respond to pro-adipogenic signals. Furthermore, the presence or absence of PPARG2 may serve as a major switch between the well and de-differentiated subtypes in human LPS.

### Therapeutic targeting of the IGF1/IGF1R/PPARG signaling axis

The combination of the above single-nucleus multiome and *in vitro* analyses identified three molecular features of DDLPS: 1) chromatin inaccessibility within and loss of expression of *IGF1*, 2) the inability to transcriptionally express or upregulate *PPARG2*; and 3) a relative overexpression of *IGF1R*. Leveraging the defined IGF1/IGF1R/PPARG2 signaling deficits observed in DDLPS cells, we aimed to explore novel therapeutic strategies in cell line LPS models. First, we investigated whether *PPARG2* expression loss could be overcome by hyperactivation of PPARG1. Activated PPARG1 has increased TF binding affinity to adipose-specific downstream genes, including *PPARG2*. Others have shown that patient WDLPS cells cultured *ex vivo* with high concentrations of insulin (acting through IGF1R) and PPARG agonists had adipogenic potential(*14*); however, the effect of PPARG agonists on DDLPS cells deficient in PPARG2 are unknown. To explore this, we added rosiglitazone (a PPARG1 and PPARG2 agonist) to IGF1-DM to culture LPS6 cells or adipose derived human MSCs. We observed that the addition of rosiglitazone was not able to induce expression of *PPARG2* or increase the efficiency of *ADIPOQ* induction (Figure S12A) in DDLPS. This result suggests that hyperactivated PPARG1 alone, in the absence of appreciable PPARG2, is not sufficient to promote DDLPS cell differentiation. This finding is consistent with observed inaccessible chromatin at the *PPARG2* TSS, which may inhibit TF binding of activated PPARG1.

Given the epigenetic enrichment of GLP-1-induced regulation of insulin signaling in DDLPS, we also investigated whether GLP-1 could promote tumor cell differentiation in DDLPS through the addition of a GLP-1 agonist to IGF1-DM. Liraglutide is a long-acting GLP-1 agonist thought to promote adipocyte differentiation by propagating insulin signaling(*28, 29*). However, when we exposed DDLPS cells to liraglutide, we did not observe an increased expression of *PPARG2*. Interestingly, liraglutide exposure did result in a robust increase in *ADIPOQ* gene expression within DDLPS tumor cells relative to IGF1-DM alone, though not to the level seen in MSCs with liraglutide exposure (Figure S12B). These results suggest that terminal differentiation of DDLPS with PPARG or GLP-1 agonists is unlikely to be an effective DDLPS treatment strategy on its own, consistent with past clinical trials. Thus, the lack of preclinical efficacy for existing pro-adipogenic therapeutics to overcome IGF1 signaling dysregulation in DDLPS is consistent with the finding that *PPARG2* gene expression is not inducible in DDLPS.

Although direct targeting of PPARG or insulin/GLP-1 may not specifically impact IGF1-associated DDLPS biology, IGF1 expression loss also correlates with a relative overexpression of IGF1R in both WDLPS and DDLPS compared to normal adipose tissue (Figure 4B). IGF1R canonically acts as a plasma membrane-bound RTK and we confirmed that IGF1R localized to the plasma membrane in LPS cells (Figure 6A). Previously, IGF1R monoclonal antibodies were investigated in several phase I and II trials across multiple cancer types and demonstrated acceptable toxicity profiles(*30*–*33*). In an effort to improve the anticancer impact of monoclonal antibodies, IGF1R-ADCs have been developed(*34*), and there is an ongoing trial of an IGF1R-ADC across advanced solid tumors (W0101, NCT03316638). We therefore hypothesized that an IGF1R-targeted ADC (IGF1R-ADC) would be lethal in both WDLPS and DDLPS cells. To investigate whether an IGF1R-ADC would be cytotoxic to LPS cells, we exposed the WDLPS-derived 93T449 cell line, as well as a normal human cell line (non-transformed human aortic endothelial cells, HAECs) as a negative control, to increasing concentrations of an IGF1R-ADC conjugated to an anti-mitotic chemotherapeutic agent, monomethyl auristatin E (MMAE)(*34*). We found that the IGF1R-ADC was specifically cytotoxic to LPS cells (Figure 6B). These analyses serve as preclinical rationale for the use of IGF1R-targeted agents in LPS. Ongoing investigation of the IGF1/PPARG signaling axis in LPS may yield further opportunities for drug development.

**Figure 6.**
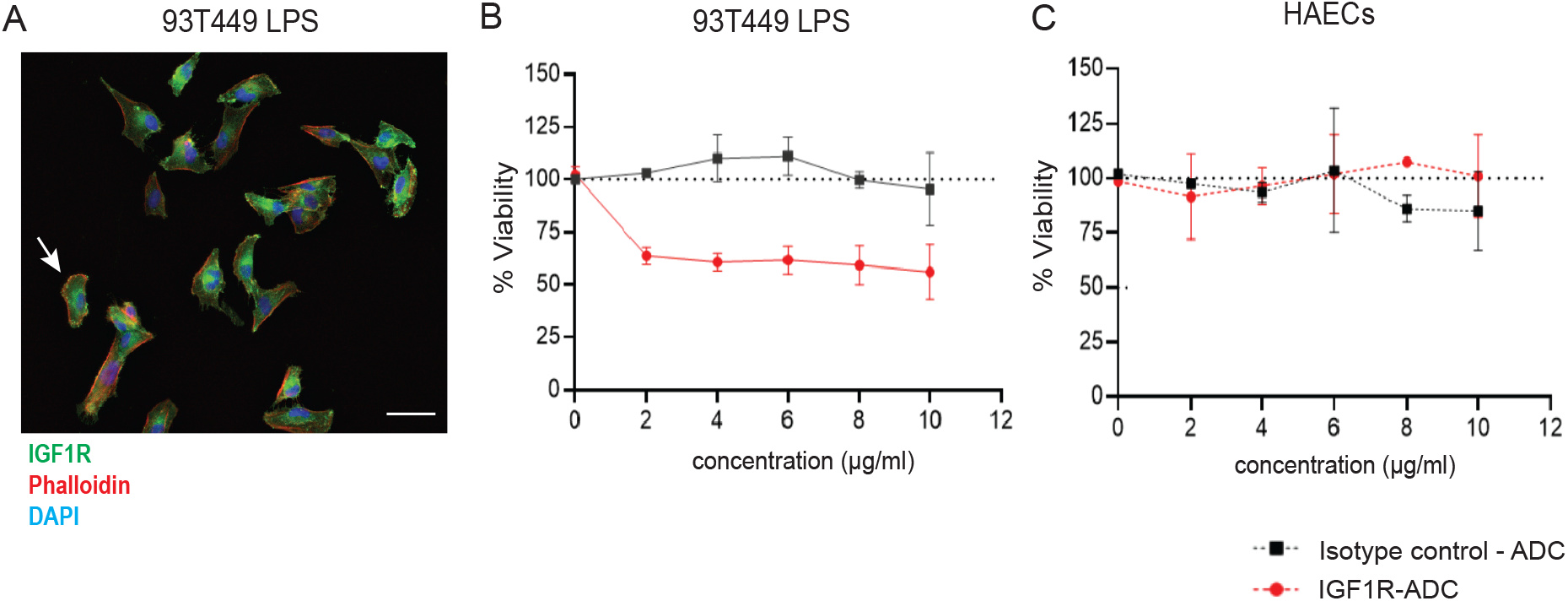
IGF1R-ADC *in vitro* cytotoxicity. (A) Immunofluorescence (IF) image of 93T449 cells illustrating membranous localization of IGF1R (green), highlighted by the white arrow. Phalloidin is shown in red and nuclear staining (DAPI) in blue. (B) 93T449 cells and (C) Human aortic endothelial cells (HAECs) were exposed to increasing concentrations (0-10 µg/ml) of anti-IGF1R or isotype control antibody conjugated to MMAE (IGF1R-ADC or Isotype control-ADC, respectively) in complete cell culture media for 72 hours. Cell viability was determined by XTT assay, setting 0 µg/ml isotype control as 100% viability.

## DISCUSSION

This study defines LPS subtype-specific molecular defects in adipocyte differentiation. We first performed single-nucleus multiome sequencing of human LPS patient tissue samples. The choice of a single-nucleus sequencing technique is a critical consideration in mesenchymal tumors and LPS specifically, as it allows for the characterization of both dense, fibrous tissues as well as larger adipocytes as opposed to previous single-cell sequencing modalities(*8, 35*). Multiome sequencing and spatial profiling from a carefully selected cohort of LPS patient samples enabled high-resolution dissection of differences in transcriptomic and epigenomic tumor cell programs and their spatial patterning. The integrated multiome sequencing analysis converged on IGF1/PPARG signaling loss as a key feature of most DDLPS cells and higher levels of *IGF1* expression in DDLPS were associated with longer overall survival. By coupling the multiome data with *in vitro* analyses of DDLPS cell lines and LPS patient tissue samples, we were able to further identify the loss of *PPARG2* gene expression as an additional defect of adipose differentiation in DDLPS. Finally, we leveraged the relative overexpression of *IGF1R* in LPS to demonstrate that IGF1R-ADCs may have clinical utility in LPS.

Our work additionally confirms findings from several other sequencing-based studies of LPS. We found transcriptomic enrichment of adipogenesis pathways in WDLPS as shown in Hirata et al.(*5*) The previously published maturity score in LPS, which predicts tumor cell differentiation and inversely correlates with *MDM2* expression(*6*), also likely encapsulates the biology of the IGF1 differentiation pathway presented in this study, thus potentially bridging MDM2 and IGF1-induced adipogenesis. Our study also provides insights regarding the general lack of response observed in early clinical trials of PPARG agonists as LPS treatments(*36*). It should be noted that many of the historical *in vitro* and clinical trial studies of PPARG agonists included numerous LPS subtypes (myxoid, pleomorphic), and we now appreciate that those driven by MDM2 amplification (e.g., WDLPS, DDLPS) represent a distinct disease entity. Our data suggest that in IGF1/PPARG2-deficient DDLPS, chromatin inaccessibility upstream of the *PPARG2* TSS prevents activated PPARG1 from binding and increasing *PPARG2* gene expression. To pursue future adipocyte differentiation-based therapies, investigating the mechanisms driving epigenetic silencing of *IGF1* and isoform-specific *PPARG* gene expression will be critical.

Leveraging the relative overexpression of *IGF1R* seen in both WDLPS and DDLPS cells compared to normal adipose tissue, we demonstrated the selective toxicity of an IGF1R-ADC in LPS cell lines. To date, there are no FDA-approved therapies for patients with WDLPS, and our preclinical data showing cytotoxicity in WDLPS-derived 93T449 cells may therefore have major clinical implications. ADCs are currently being engineered with multiple linker and payload combinations, and future formulations of an IGF1R-targeted ADC may be even more effective in LPS.

Additionally, the differential chromatin accessibility around genes involved in the GLP-1 pathway and the observed modest effect of liraglutide on markers of adipose differentiation *in vitro* underscores the need for understanding the role of GLP-1 in adipocytes, MSC differentiation, and sarcomagenesis. The exact mechanisms through which GLP-1 influences cellular processes in adipocytes remains unknown, although our study identifies a potential new role for this process in DDLPS specifically that warrants further investigation.

While we have uncovered LPS subtype-specific aberrations in adipocyte differentiation with the potential for clinical translation, our study has several limitations. First, despite consistent multiome sequencing library preparation and computational analysis across the cohort, sample-specific batch effects cannot be mitigated entirely. Additionally, the number of high-quality nuclei for analysis within the snRNA-seq and snATAC-seq from the multiome data was unbalanced. We preferentially used the highest-quality nuclei for both assays at the expense of fewer analyzable nuclei within the snATAC-seq assay. Finally, we are limited by a lack of bona fide WDLPS human cell lines with which to further investigate adipose differentiation pathways *in vitro*.

In summary, we characterized the transcriptional, epigenetic, and spatial features of WDLPS and DDLPS patient tissue samples, identified deficits in IGF1 and PPARG2 as a characteristic of most DDLPS cells that may contribute to dysfunctional adipose differentiation, and illustrated that targeting IGF1R may serve as a novel therapeutic strategy for LPS patients. Further investigation into the upstream epigenetic regulators of *IGF1* and isoform-specific *PPARG* gene expression may yield additional opportunities to improve LPS patient outcomes.

## AUTHOR CONTRIBUTIONS

Conceptualization: EMP, EMVA; Data generation and curation: EMP, CPR, JLH, ART, AN, JP, SG, GDD, ES; Analysis: EMP; Funding acquisition: GDD, NLS, EMP, EMVA; Experimental design: EMP, AEG, MJK; Experimental execution: EMP, AEG; Project administration: EMP, JP, NS, ES; Resources: NS, MK, JP, EMVA; Supervision: EMVA, ART, AN, GDD, NLS; Validation: EMP; Data visualization: EMP; Writing, original draft: EMP, EMVA, AEG, SYC; Writing, editing and review: JP, SLH, LV, JF, KB, BMT, ES, YJK, ART, AN, CPR, PM, NLS, SG, GDD

## METHODS

### Sample collection and nuclear isolation

Human tissue samples were collected after written informed consent obtained under the Institutional Review Board-approved Dana-Farber Cancer Institute research protocol #21-468. Independent pathological review of each patient sample was performed by Dr. Jason Hornick (Director of Surgical Pathology and Immunohistochemistry, Department of Pathology, Brigham and Women’s Hospital).

Resection specimens of human LPS tumor or adipose tissue were collected and snap-frozen in liquid nitrogen. Nuclei were isolated in accordance with 10x Genomic Chromium Next GEM Single Cell Multiome ATAC+ Gene Expression User Guide (Rev E) by first mincing the tissue with spring scissors and triturating with a pipette for a total of 10 min in Tris-Salt-Tween (0.03% Tween) extraction buffer on ice(*35*). The resulting nuclei suspension was filtered through a 30 um MACS SmartStrainer. Filtered nuclei were centrifuged (500 x g, 5 min, 4C) and the pellet was resuspended and permeabilized in a lysis buffer on ice for 2 minutes. Nuclei were then washed, centrifuged, and resuspended in Diluted Nuclei Buffer. Nuclei were stained with DAPI (Invitrogen ProLong Gold Antifade Mountant with DAPI) and manually counted on a hemocytometer. A maximum of 16,000 nuclei per sample were loaded onto a channel of the Chromium Next GEM Chip J for use with the 10x Chromium Controller. Following transposition, snRNA-seq and snATAC-seq libraries were prepared following the 10x Genomics Multiome protocol. Library metrics were analyzed with the Agilent Bioanalyzer and sequenced per the specifications in the 10x Genomic Multiome User Guide. All libraries were sequenced by the Molecular Biology Core Facilities at Dana-Farber Cancer Institute using an Illumina NovaSeq 6000 sequencer.

### snRNA-seq data preprocessing

10x Cell Ranger ARC v2.0.0 (Cell Ranger v6.1.1) was used to demultiplex multiome sequencing data, process barcodes, align sequencing reads to genome GRCh38, and perform UMI counting. Non-duplicate reads that mapped confidently to the genome were counted to form a gene-barcode matrix. To capture as many high-quality nuclei from each snRNA/ATAC-seq assay individually, the cellranger ARC output raw feature matrix was carried forward for downstream pre-processing and analysis. First, quality control and filtering of cell nuclei were performed for each snRNA-seq and snATAC-seq assays separately per patient sample.

Within the snRNA-seq data, Cellbender v0.2.2(*37*) was used to remove barcodes representing empty droplets. The learning rate, epochs, and total droplets included were adjusted per sample to ensure an appropriate training curve. Scrublet v0.1(*38*) was used to simulate droplets containing doublets in the snRNA-seq data. Expected doublet rate was calculated based on number of barcodes per sample after filtering for empty droplets (https://demultiplexing-doublet-detecting-docs.readthedocs.io/en/v0.0.4/test.html). A manual threshold was set for each sample bisecting a bimodal distribution of cells (presumed singlets and doublets). Droplets predicted to be doublets were removed from the gene-barcode matrix of the snRNA-seq data. Scanpy v1.9.3(*39*) was used for further filtering of low-quality nuclei, where nuclei with fewer than 500 UMI, fewer than 200 genes, or more than 5% of mitochondrial transcripts were removed. After, gene expression values were log-normalized, filtered for highly variable genes, and scaled. Linear dimensionality reduction with principal component analysis (PCA) was performed and a KNN graph was calculated (n_neighbors = 15), before non-linear dimensionality reduction and visualization with Uniform Manifold Approximation and Projection (UMAP). Unsupervised Leiden clustering was performed using the first 40 principal components (PCs) with a resolution between 0.2-0.4. Differential gene expression between each Leiden cluster and the aggregate of all other clusters was performed with a two-sided Wilcoxon rank-sum test and Bonferroni correction. The resulting DEGs were used to broadly annotate cluster cell types. Inferred copy number alteration (CNA) analysis was performed using both the inferCNV (https://github.com/broadinstitute/inferCNV) and SCEVAN(*40*) algorithms, using immune cells as the reference cell type. From this, malignant cells were identified by the presence of predicted CNA amplification at chromosome 12q, as well as differential expression of known marker genes.

After filtering nuclei and designating broad cell type annotations per individual patient sample snRNA-seq data, raw gene expression counts from each individual patient sample were then merged. After merging, genes detected in less than three cells and nuclei barcodes with more than 8,000 genes were additionally filtered. Expression values were then log-normalized and scaled across all merged patient nuclei. As each sample was processed individually, Harmony_py v0.0.6(*41*) was used to mitigate potential batch effects prior to performing dimensionality reduction and unsupervised clustering. After batch effect correction with Harmony_py, PCA was performed on the first 50 PCs, KNN graph calculated (n_neighbors = 15), followed by Leiden clustering at a resolution of 0.4 and UMAP visualization. Further downstream analysis was performed on either the raw or log-normalized and scaled expression values as specified below.

### Differential gene expression, cNMF, and gene set enrichment analysis

To further evaluate the LPS tumor cells, the 80,881 nuclei annotated as tumor were isolated, re-integrated, and clustered with 40 PCs at a resolution of 0.4 as detailed above. Differential gene expression analysis comparing WDLPS and DDLPS tumor cells was performed on the log-normalized expression values. Gene set enrichment of Hallmark 2020 and Gene Ontology Biological Processes gene sets from MSigDB(*42*) was evaluated with BEANIE(*18*) using the genes derived from differential gene expression analysis between WDLPS and DDLPS cells.

To identify gene activity programs, cNMF analysis(*19*) was performed on the LPS tumor cells. Raw counts were used as input to the cNMF algorithm with parameters of 200 iterations, 5000 genes, a seed of 333, and a k range of 5-15. Balancing the calculated stability and error rate, cNMF usage programs identified at k = 7 were selected for further investigation. Gene set overlap(*43*) analysis (FDR < 0.05) with MSigDB Hallmark, Gene Ontology Biological Processes, Kegg legacy, and medicus gene sets(*42*) was performed using the genes derived from the cNMF usage program gene lists (top 200 genes).

### snATAC-seq data preprocessing

FASTQ files were input to Cell Ranger ARC v2.0.0 for read filtering and alignment (GRCh38). For each sample, a peak-barcode matrix was constructed by calling peaks on the sample fragment file using MACS3 within Signac v1.8.0(*44, 45*). The matrix was then used to create a chromatin assay and Seurat object (Seurat v4.3.0) with CreateChromatinAssay and CreateSeuratObject functions at default thresholds. Doublets were removed using scDblFinder v1.12.0(*46*) with default parameters. The following quality control metrics were used to filter out barcodes that did not meet the following criteria: peak region fragments > 1000, FRiP > 0, TSS > 2, and blacklist fractions and nucleosome signals that were < 3 times the mean absolute deviation from the median. Nuclei that passed quality control underwent normalization with term frequency-inverse document frequency (TF-IDF) and linear dimensionality reduction with singular value decomposition (SVD) to produce latent semantic indexing components (LSI). Non-linear dimensionality reduction with UMAP was performed, nearest neighbors were computed, and unsupervised clustering was performed at a resolution between 0.1-0.3. Nuclei clusters were broadly annotated by cell type with information from the following analysis: differential gene activity (GeneActivity function, FindAllMarkers with latent.var = peak_region_fragments, test.use = LR), marker gene activity, and by inferring CNAs with CopyscAT v0.40(*47*).

To perform analysis on a cohort-level object across all patient samples, individual patient sample-level peak sets were combined to create a common peak set. Common peaks across all patient samples were quantified per individual patient sample using the sample-level fragment file. The resulting peak-by-cell matrices per individual patient sample were merged to generate the aggregated cohort-level object. Quality control was performed on the cohort with the same metrics described above followed by normalization, dimensionality reduction, clustering, and UMAP visualization (dimensions 2-8 and resolution of 0.1). Non-malignant cell types clustered together so integration was not performed. Peaks were re-called by cell type and served as the final peak set for analysis. With the final peak set, we created the final ATAC object by constructing a cohort-level count matrix, performing normalization, dimensionality reduction, and clustering.

### Differential peak accessibility and genomic enrichment analysis

Tumor-adipose differential accessibility analysis was performed by subsetting the cohort-level ATAC object to cells labeled “adipocyte,” “WDLPS,” and “DDLPS,” before normalization, dimensionality reduction, and clustering, as above. Adipocytes were included in this analysis as a positive control for the biological relevance of ATAC data given the relative low number of cells compared to snRNA-seq analysis. Differential accessibility was calculated with Signac FindAllMarkers using a logistic regression adjusting for sequencing depth (nCount_ATAC) as a latent variable. Differentially accessible peaks with an adjusted p-value of < 0.05 (Bonferroni-Hochberg correction) and log2 fold change of at least 1 were considered for downstream analysis. BED files from the differentially accessible peaks of each group were created and coordinates were lifted over to the hg19 reference genome using the liftOver function from rtracklayer v1.58.0(*48*) to input as the “Test regions” with GREAT v3.0.0(*49*) with default parameters. Coverage plots for peaks overlying a gene body were generated with Signac’s CoveragePlot function. Differentially accessible peaks specific to *IGF1* were calculated by first identifying peaks in the tumor-adipose object that overlapped with coordinates of the *IGF1* gene body using the findOverlaps function. Next, the tumor-adipose count matrix was subsetted to include only those peaks that overlap *IGF1*. Finally, differential accessibility across *IGF1* in WDLPS, DDLPS, and adipocytes was calculated with FindMarkers (where ident.1 = WDLPS and ident.2 = DDLPS) or FindAllMarkers functions. Candidate response element genomic coordinates were derived from the UCSC Genome Browser (hg38)(*50*). Similar analysis was carried out for peaks within the *PPARG* gene body.

### Bulk transcriptomic analysis, signature scoring, and survival analysis

Raw bulk RNA-seq data of LPS and normal adipose patient samples published in Bevill et al(*6*) was processed using Scanpy v1.9.3 with CPM normalization and log2 transformation. Log-normalized (TPM) cell line bulk RNA-seq data was obtained from the Broad Institute’s Cancer Cell Line Encyclopedia(*51*). The IGF1 autocrine signature score was calculated by summing the expression values of the positively regulated genes *(IGF1, PPARG, FABP4, LPL*) and subtracting the expression of *IGF1R*. Statistical significance was assessed by the Wilcoxon rank-sum test.

For survival analysis, transcriptomic data from patients with a primary diagnosis of “dedifferentiated liposarcoma” or “liposarcoma, well differentiated” were downloaded from The Cancer Genome Atlas (TCGA) Research Network: https://www.cancer.gov/tcga. Raw expression values underwent variance stabilizing transformation using the DESeq2 package v1.42.0(*52*). The median expression value of *IGF1* expression in the TCGA cohort was calculated (median value = 8.1655) and used as a cutoff to divide the cohort into “IGF1 high” and “IGF1 low” groups. A Kaplan-Meier curve was used to visualize the overall survival of each group using the survfit function and survival probabilities were calculated with the survdiff (rho = 0) function of the survival package v3.7_0(*53*).

### Spatial transcriptomic sample preparation and library construction

Spatial gene expression profiling of FFPE tissues was performed with the 10x Genomics Visium CytAssist Spatial Gene Expression (Rev E). Tissue sections of 10-µm thickness were placed on a slide, and H
&E staining was performed by the Brigham and Women’s Hospital Pathology Department. Additional scrolls of 10-µm thickness were collected and stored at 4^0^C. RNA quality was assessed using RNA extracted from scrolls with the Qiagen RNeasy FFPE kit and integrity measured by DV200 value with the Agilent 4200 TapeStation with RNA High Sensitivity Screen Tape. FFPE H&E slides were imaged according to the Visium Imaging Guidelines Technical Note. The sample was then de-crosslinked and de-stained all in accordance with the 10x Genomics Visium CytAssist Spatial Tissue Preparation Guide (CG00518 Rev C). Library construction and sequencing of the 6.5 mm x 6.5 mm tissue section was completed according to the Visium CytAssist Spatial Gene Expression User Guide (CB000495, Rev E) using Visum Human Transcriptome probe set v2.0 (18,536 genes targeted by 54,518 probes). Libraries were normalized and pooled for sequencing on Illumina NextSeq 150 flow cells (Illumina, Inc., San Diego, CA, USA).

### Spatial transcriptomic data processing and signature scoring

Space Ranger v2.1.1 was used to demultiplex sequencing data, process barcodes, input slide images, align sequencing reads to genome GRCh38, and perform UMI counting. Spatial transcriptome sequencing data was analyzed using Squidpy v1.2.3(*54*). Filtering for high quality barcodes (spots) was based on the following metrics: counts >1,000 and < 30,000, mitochondrial read percentage < 20%, and genes filtered based on their presence in 10 cells or more. Counts were normalized using scTransform v0.01(*55*) for Python. Normalized values underwent dimensionality reduction, UMAP visualization, and clustering at a resolution of 0.8. The IGF1 autocrine signature score was applied to the spatial data and visualized. Additionally, cNMF programs were assessed as a signature score by summing the expression of the top 50 genes in each program. Comparison of spatial distribution of the two cNMF signature scores was assessed by designating each spot as belonging to the cNMF program for which it had the highest score.

### Cell lines and differentiation

LPS6 human DDLPS-derived cells were a generous gift from the Shapiro Lab (DFCI) and were maintained in RPMI-1640 medium, supplemented with 10% FBS, 1% glutaMaX, 1,000 U/ml penicillin and 1 mg/ml streptomycin. The 93T449 WDLPS-derived cell line and adipose-derived human mesenchymal stem cells (MSCs) were purchased from ATCC and cultured according to the manufacturer’s instructions. Human Mesenchymal Stem Cells (MSC) were maintained in mesenchymal stem cell basal medium supplemented with the HMSC growth kit-low serum (ATCC). Human Aortic Endothelial Cells (HAOEC) were purchased from Milipore Sigma and maintained in endothelial cell growth medium MV with supplement mix, containing 10% FBS (PromoCell). Cells were grown at 37^0^C in a humidified atmosphere with 5% CO2. Media was replaced every two to four days and cells passaged at approximately 70-80% confluency. For adipogenic differentiation, LPS cell lines were seeded at 30-50% confluency and treated with 10% charcoal stripped FBS with a cocktail of dexamethasone (Dex 1 mM), indomethacin (IDM, 50 mM), 3-isobutyl-1-methylxanthine (IBMX, 0.5 mM), and human recombinant IGF-1 (Miltenyi Biotec, 200 nM). Media and differentiation cocktail was replaced every three days. Differentiation induction measured at day 3.

### Gene Expression Analysis/Real-Time PCR

Total RNA from cell lines was extracted using the Qiagen RNeasy Mini Kit following the manufacturer’s instructions. The quantity of RNA was measured using a NanoDrop 8000 spectrophotometer. For cDNA synthesis, 0.5 - 1 µg of total RNA was reverse transcribed using the High-capacity cDNA RT kit (Applied Biosystems) according to the manufacturer’s instruction and performed in a PTC Tempo Thermal Cycler (BioRad). Real-time quantitative PCR (QPCR) was performed using the 7500 Fast Real-time PCR system and Powerup SYBR Master mix (Applied Biosystems). Each 10 µl reaction contained master mix, 1 µl cDNA template, and 300 nM primer concentration. Primers were designed using NCBI Primer Blast tool and oligos were synthesized by Integrated DNA Technologies (IDT). Primers for target genes (*PPARG, ADIPOQ*, and reference gene (*β-actin*) are listed in Table 1. The relative expression levels of the target genes were calculated using the ΔΔCt method with *β-actin* as the internal control. All reactions were performed in duplicate, and data are presented as mean +/-SD.

**Table 1.**
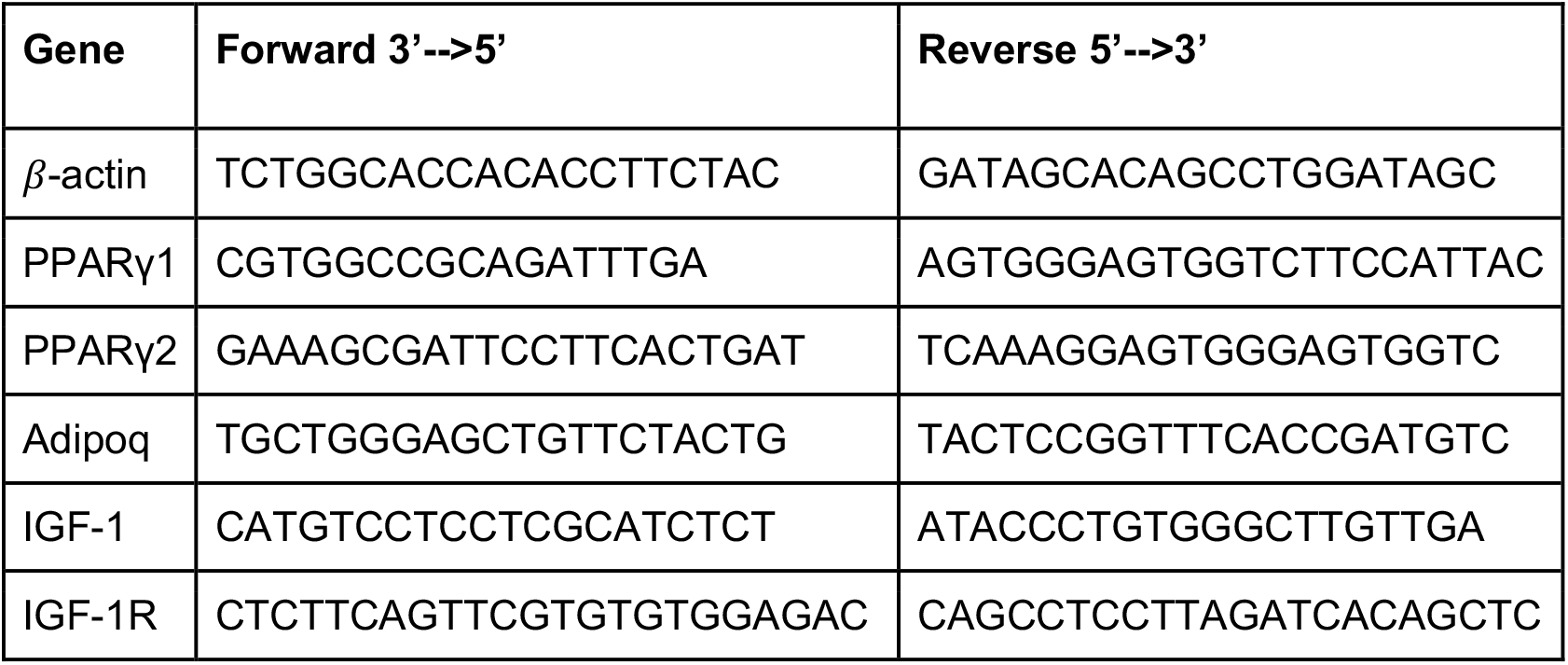
Primers used in QPCR.

### Immunostaining of liposarcoma cells

Cells were cultured on 20 mm round coverslips for 1–2 days, rinsed with PBS, and fixed with 4% formaldehyde freshly prepared in PBS for 15 minutes at room temperature. Following fixation, the cells were rinsed three times with PBS. For permeabilization and blocking, cells were incubated for 60 minutes at room temperature in blocking buffer composed of 1X PBS, 5% normal serum (CST #5425), and 0.3% Triton X-100. Coverslips were then incubated overnight at 4°C with either primary antibody IGF-1Rβ (D23H3) (1:100 dilution, #9750, Cell Signaling Technologies) or dilution buffer alone (PBS with 1% BSA and 0.3% Triton X-100) as a negative control. The next day, cells were rinsed with PBS and incubated with secondary antibody (anti-rabbit IgG (H+L), F(ab’)2 Fragment, Alexa Fluor® 488, #4412, CST) for 2 hours in the dark at room temperature. Afterward, coverslips were washed three times with PBS and stained with Alexa Fluor® 555 Phalloidin (CST) for 15 minutes at room temperature, followed by a final PBS rinse. Coverslips were mounted onto microscope slides using ProLong Gold Antifade Reagent with DAPI (#8961, CST). Fluorescence imaging was performed using a Leica THUNDER Imager, and final image processing was completed with ImageJ software.

### Antibody drug conjugate cell viability assays

IGF1R-targeted antibody drug conjugate (IGF1R-ADC) and IgG1 isotype control ADC were constructed as previously described (Akla et al.(*34*)) and purchased from Creative Biolabs (Shirley, NY). Specifically, lonigutamab antibody was linked to mc-MMAE (monomethyl auristain E) with a drug-to-antibody ratio (DAR) of 4. Human IgG1 isotype control antibody was also linked to mc-MMAE with a DAR of 4. Assessment of cell viability was performed using CyQUANT XTT Cell Viability Assay™ (Invitrogen) following the manufacturer’s instructions. Briefly, cells were seeded in 96-well plates and allowed to adhere 16-24 hours prior to treatment. After 72 hours of treatment, cells were incubated with XTT working solution for 4 hours and absorbance was measured at 450 nm and 660 nm using the CLARIOstar Microplate reader.

### Statistical analysis

Graphical data was expressed as the mean +/-standard deviation and statistics were calculated using GraphPad Prism 10 software. Statistical tests are specified in figure legends.

## ACKNOWLEDGEMENTS

We thank the patients and for their participation in this study. We also thank Bruce Spiegelman for his expertise and guidance on adipogenesis and experimental design. We thank the Shapiro Lab for their generosity in supplying LPS cell lines. We thank Jamie Brett and the Haigis Lab for supplying additional LPS patient samples, Shannon Coy and the Santagata Lab for providing FFPE transition zone slides, and Kathleen Kee for her assistance in locating and organizing patient samples. The authors would like to thank the Center for Cancer Genomics at Dana-Farber Cancer Institute and the Single Cell Core at Harvard Medical School, Boston, MA for performing the single cell RNA-Seq library preparation. This project was supported by the David Liposarcoma Research Initiative via the Rossy Foundation Fund at Myraid Canada, the Doris Duke Charitable Foundation Physician Scientist Fellowship, the 2024 AACR-QuadW Foundation Sarcoma Research Fellowship in Memory of Willie Tichenor #1259709. This project was also supported by a Conquer Cancer Young Investigator Award. Any opinions, findings, and conclusions expressed in this material are those of the author(s) and do not necessarily reflect those of the American Society of Clinical Oncology® or Conquer Cancer®. Additional support for this project from R01CA278980, R37CA222574, R50CA265182, T32GM008313, the Parker Institute for Cancer Immunotherapy, The Ambrose Monell Foundation, and the National Science Foundation GRFP DGE1144152. GDD would like to acknowledge the Dr. Miriam and Sheldon G. Adelson Medical Research Foundation Ubiquitin Pathway Program of Needham, MA and the Ludwig Center at Harvard in Boston, MA for support.

