## Supplemental Figures for "Epigenetic dysregulation of metabolic programs mediates liposarcoma cell plasticity"

**Affiliations:**
[1]Department of Medical Oncology, Dana-Farber Cancer Institute, Boston, MA 02115, USA
[2]Broad Institute of Harvard and MIT, Cambridge, MA 02142, USA
[3]Sarcoma Center, Dana-Farber Cancer Institute, Boston, MA 02115, USA
[4]David Liposarcoma Research Initiative, Dana-Farber Cancer Institute, Boston, MA 02115, USA
[5]Harvard Division of Medical Sciences PhD Program in Biological and Biomedical Sciences, Boston, MA 02115, USA
[6]Harvard-MIT MD-PhD Program, Harvard Medical School, Boston, MA, 02115, USA
[7]Harvard Medical School, Boston, MA, 02115, USA
[8]Breast Oncology Program, Dana-Farber Brigham Cancer Center, Boston, MA, 02115, USA
[9]Center for Cancer Genomics, Dana-Farber Cancer Institute, Boston, MA, 02115, USA
[10]Department of Radiation Oncology, Massachusetts General Hospital, Boston, MA, 02114, USA
[11]Department of Surgery, Brigham and Women’s Hospital, Boston, MA, 02115, USA
[12]Department of Pathology, Brigham and Women’s Hospital, Boston, MA, 02115, USA
[13]Ludwig Center at Harvard, Boston, MA, 02115, USA
[14]Parker Institute for Cancer Immunotherapy, Dana-Farber Cancer Institute, Boston, MA, 02115, USA

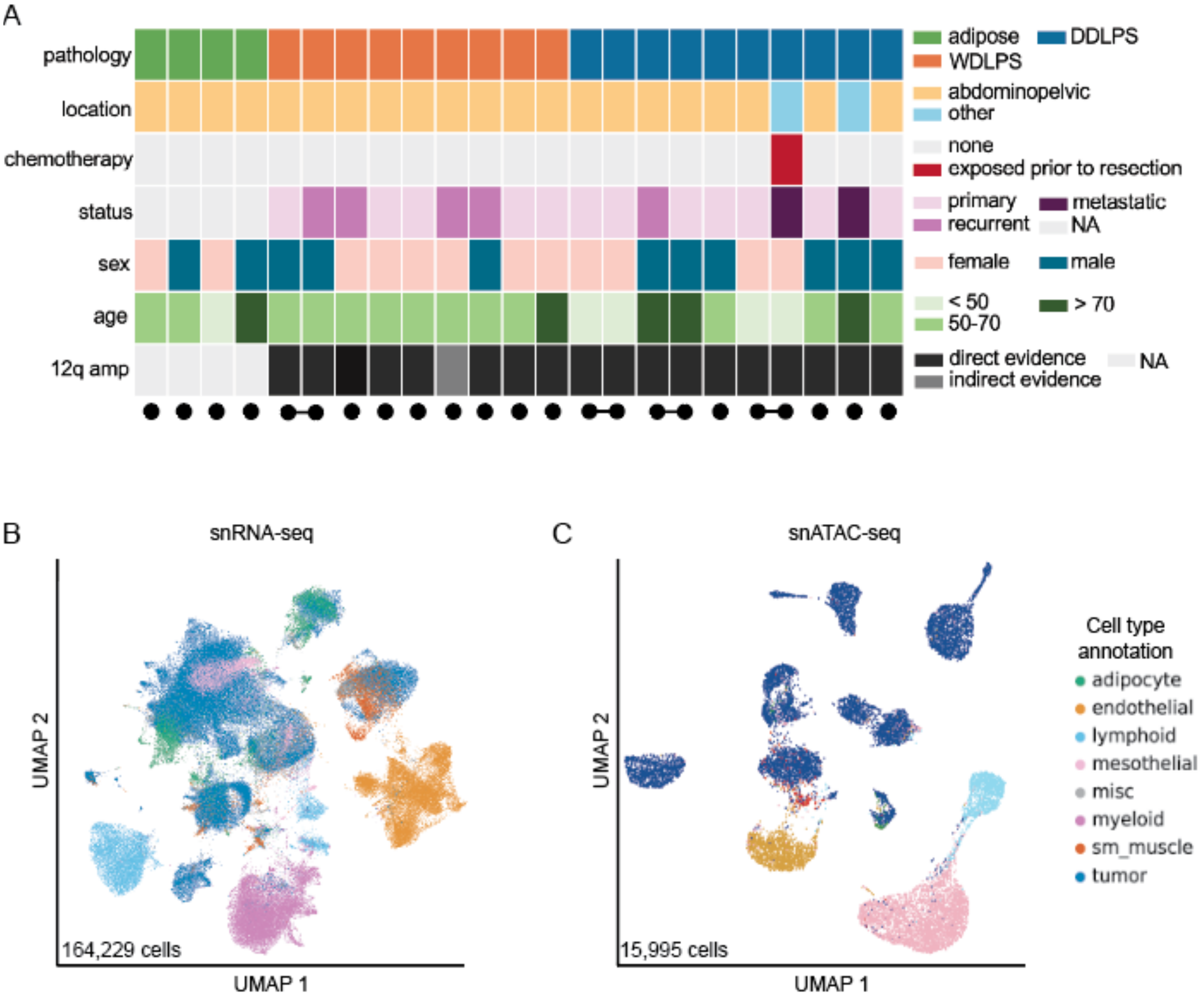


**Figure S1. Multiome analysis of LPS patient samples.** (A) Summary of clinical features at the time of patient sample resection. DDLPS, dedifferentiated liposarcoma; WDLPS, well differentiated liposarcoma; RT, radiation therapy; direct evidence refers to elevated levels of MDM2 (located on chr12q) within the sample and indirect signifies the presence of ringed chromosomes in the sample. On the bottom, each patient is represented by a solid black dot. Patients with multiple samples are linked with a line between them. (B) Uniform manifold approximation and projection (UMAP) of cells captured across all samples via single nucleus RNA-sequencing (snRNA-seq). C. UMAP of cells captured across all samples via snATAC-seq. Cell type annotation legend applicable to B and C.

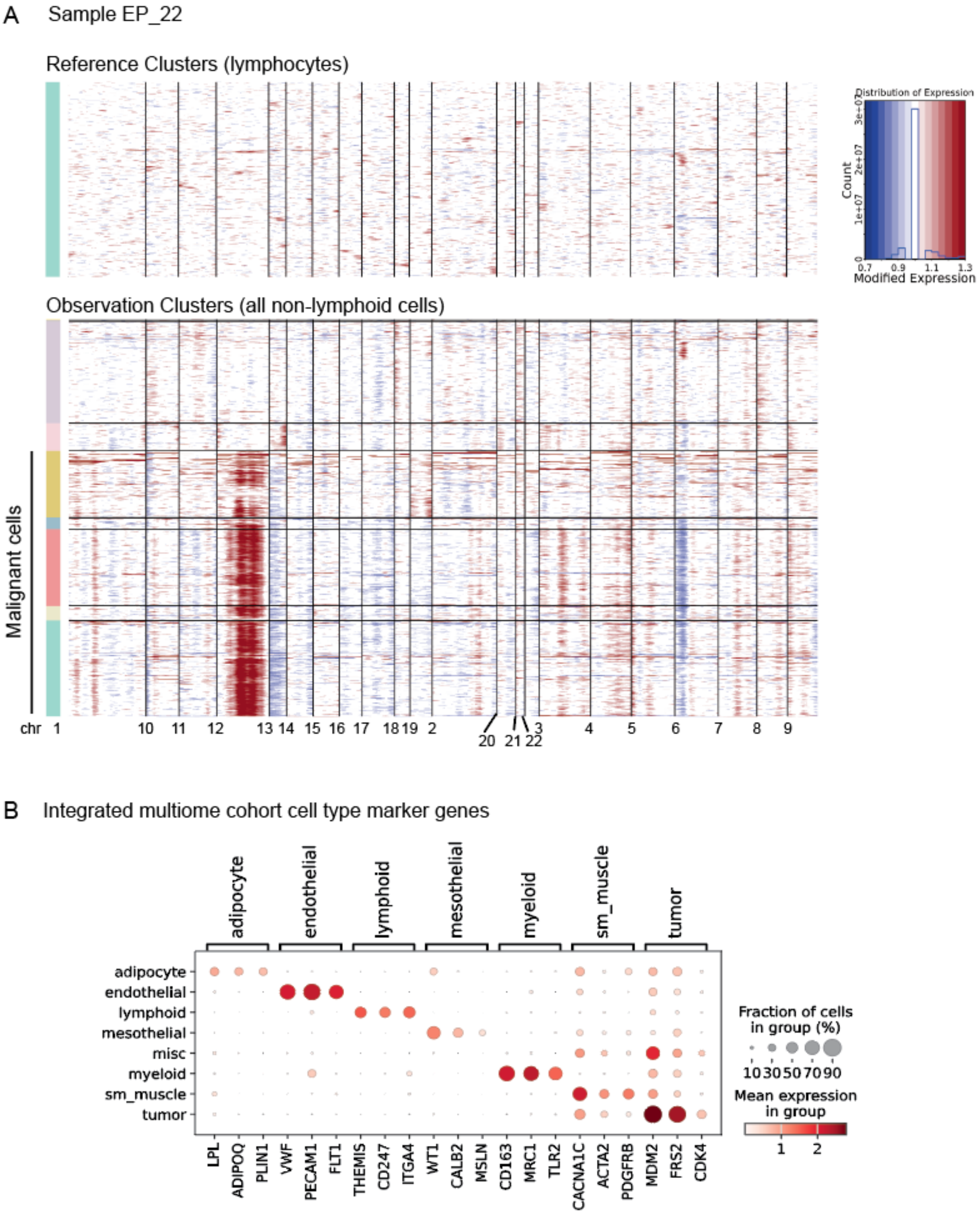
A
Sample EP_22
Reference Clusters (lymphocytes)
Distribution of Expression
Count
Modified Expression
Observation Clusters (all non-lymphoid cells)
Malignant cells
chr 1 10 11 12 13 14 15 16 17 18 19 2 20 21 22 3 4 5 6 7 8 9
B
Integrated multiome cohort cell type marker genes
adipocyte
endothelial
lymphoid
mesothelial
misc
myeloid
sm_muscle
tumor
LPL ADIPOQ PLIN1 VWF PECAM1 FLT1 THEMIS CD247 ITGA4 WT1 CALB2 MSLN CD163 MRC1 TLR2 CACNA1C ACTA2 PDGFRB MDM2 FRS2 CDK4
Fraction of cells in group (%)
10 30 50 70 90
Mean expression in group
1 2

**Figure S2. snRNA-seq derived cell type annotation.** Cell type annotation began at the sample level, where broad labels were applied to each cluster based on differentially expressed genes. Copy number analysis was performed on a per-sample basis. (A) Example InferCNV output for patient sample EP_22, where red indicates amplification and blue a deletion relative to indicated reference cells. Cell clusters with amplification of chr 12 as well as high expression of genes on the amplicon (e.g. MDM2, FRS2) were annotated as ‘tumor’. Once each sample was broadly annotated, samples were integrated and re-analyzed for more granular cell type annotation based on marker gene expression. (B) Dot plot of relative marker gene expression for indicated cell types across all cells in the snRNA-seq multiome cohort that passed QC (Methods). The label ‘tumor’ includes both WDLPS and DDLPS tumor cells. sm_muscle = smooth muscle cells.

A Sample EP_1

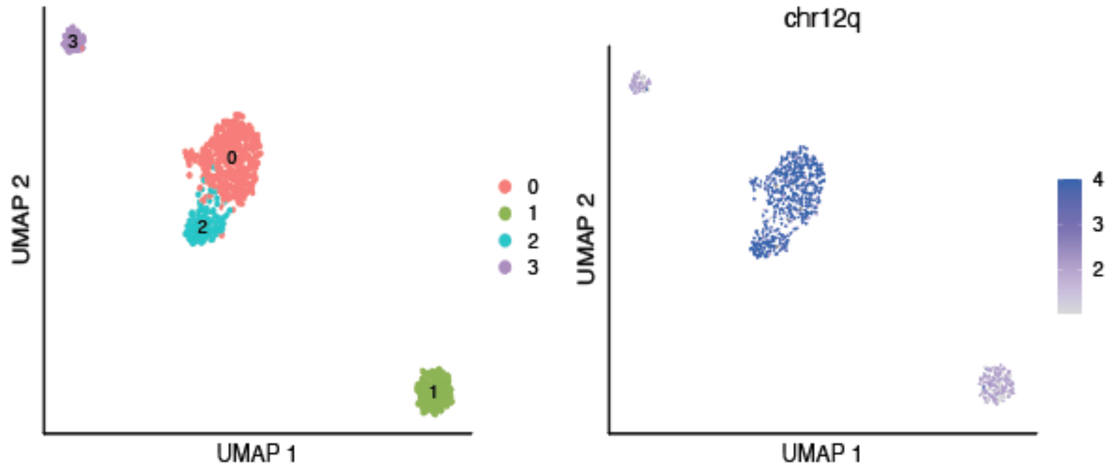

B

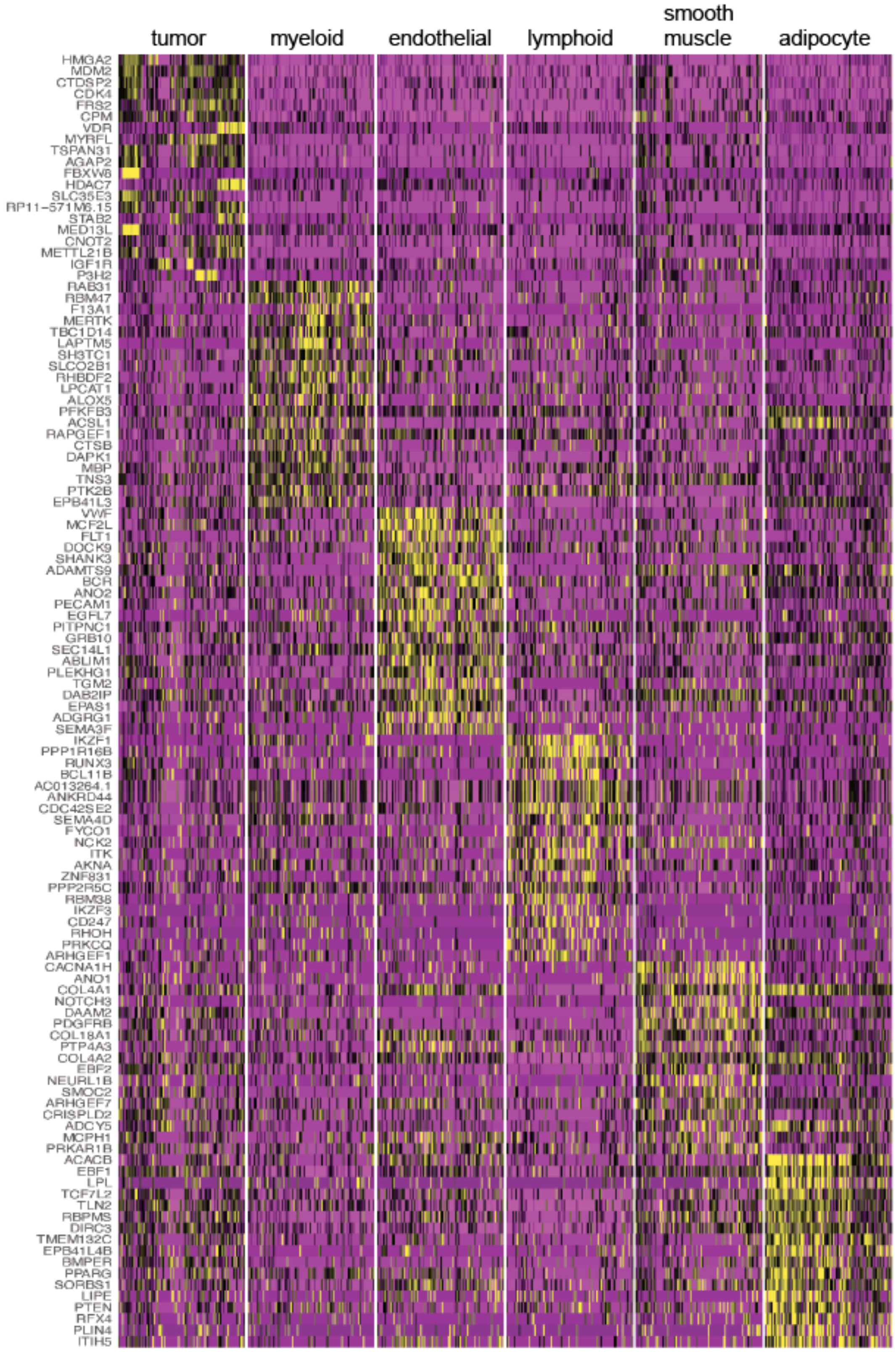

tumor
myeloid
endothelial
lymphoid
smooth muscle
adipocyte
HMGA2
MDM2
CTDSP2
CDK4
FRS2
CPM
VDR
MYRFL
TSPAN31
AGAP2
FBXW8
HDAC7
SLC35E3
RP11-571M6.15
STAB2
MED13L
CNOT2
METTL21B
IGF1R
P3H2
RAB31
RBM47
F13A1
MERTK
TBC1D14
LAPTM5
SH3TC1
SLCO2B1
RHBDF2
LPCAT1
ALOX5
PFKFB3
ACSL1
RAPGEF1
CTSB
DAPK1
MBP
TNS3
PTK2B
EPB41L3
VWF
MCF2L
FLT1
DOCK9
SHANK3
ADAMTS9
BCR
ANO2
PECAM1
EGFL7
PITPNC1
GRB10
SEC14L1
ABLIM1
PLEKHG1
TGM2
DAB2IP
EPAS1
ADGRG1
SEMA3F
IKZF1
PPP1R16B
RUNX3
BCL11B
AC013264.1
ANKRD44
CDC42SE2
SEMA4D
FYCO1
NCK2
ITK
AKNA
ZNF831
PPP2R5C
RBM38
IKZF3
CD247
RHOH
PRKCQ
ARHGEF1
CACNA1H
ANO1
COL4A1
NOTCH3
DAAM2
PDGFRB
COL18A1
PTP4A3
COL4A2
EBF2
NEURL1B
SMOC2
ARHGEF7
CRISPLD2
ADCY5
MCPH1
PRKAR1B
ACACB
EBF1
LPL
TCF7L2
TLN2
RBPMS
DIRC3
TMEM132C
EPB41L4B
BMPER
PPARG
SORBS1
LIPE
PTEN
RFX4
PLIN4
ITIH5

**Figure S3. snATAC-seq derived cell type annotation.** Cells were first broadly annotated using gene activity (a proxy for expression) for marker genes as well as copy number analysis to identify tumor cells. A. Sample level example of cell clusters (left) and output from CopyScAT, with inferred amplification of chr12q in clusters 0 and 1. B. Marker gene activity per major cell type in all cells of the snATAC-seq multiome cohort that passed QC for ATAC-seq analysis.

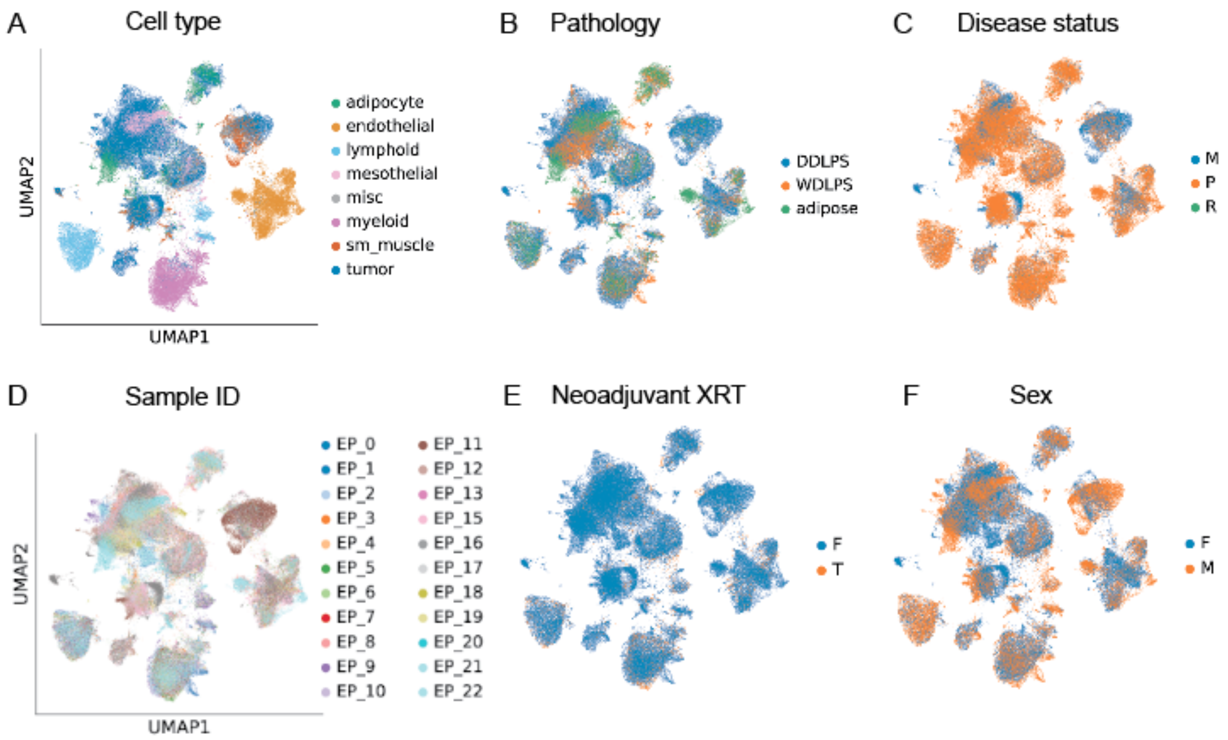


**Figure S4. Clinical cohort characteristics mapped onto snRNA-seq derived UMAP of all cell types.** Cell clusters by cell type (A), pathology (B), disease status (C), sample ID (D), exposure to neoadjuvant radiation (XRT) prior to resection (E), and patient sex (F). Disease status; metastatic = M, primary abdominopelvic = P, recurrent abdominopelvic = R. Neoadjuvant XRT; no exposure = False (F), exposure = True (T). Sex; Female = F, Male = M.

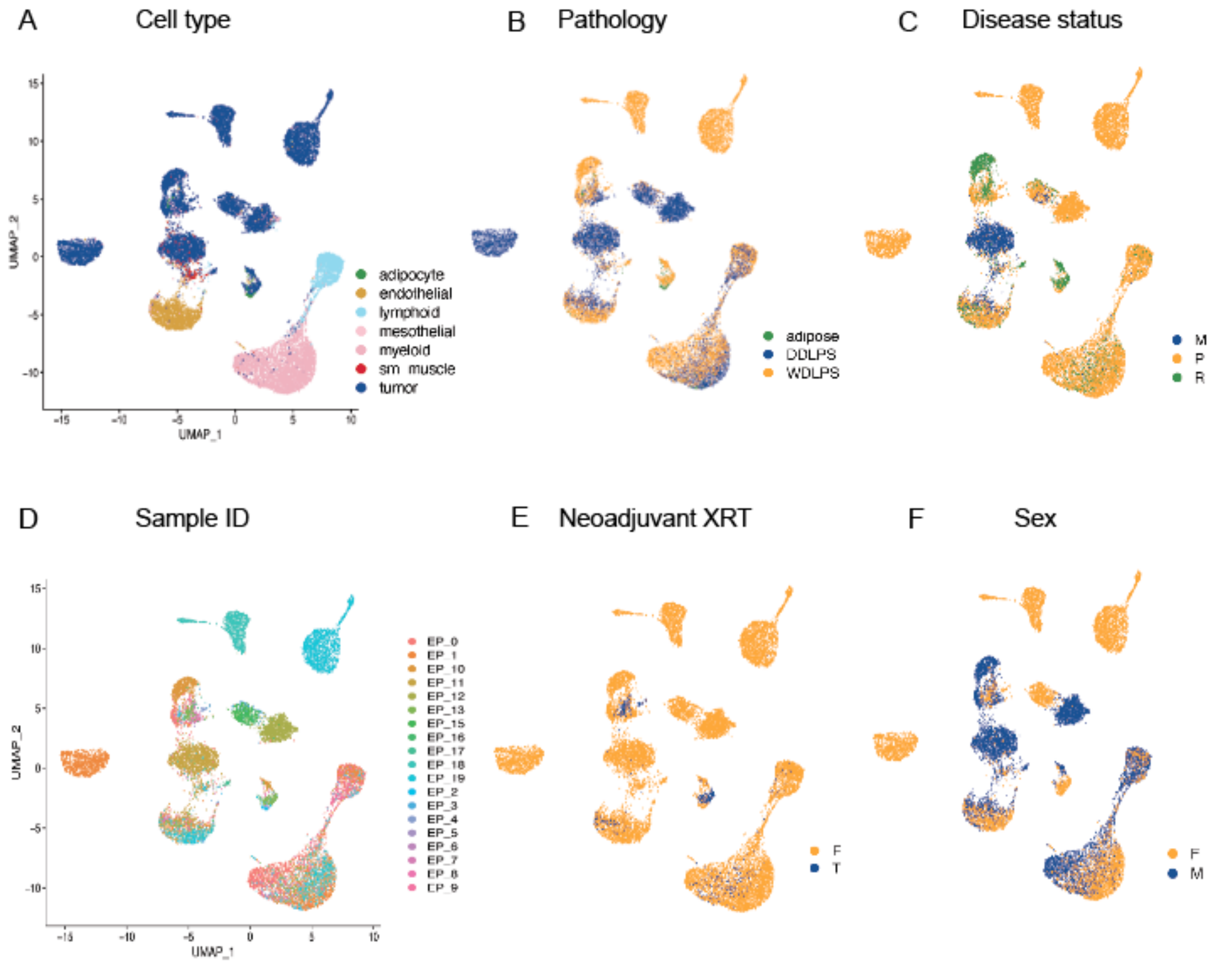


**Figure S5. Clinical cohort characteristics mapped onto snATAC-seq derived UMAP of all cell types.** Cell clusters by cell type (A), pathology (B), disease status (C), sample ID (D), exposure to neoadjuvant radiation (XRT) prior to resection €, and patient sex (F). Disease status; metastatic = M, primary abdominopelvic = P, recurrent abdominopelvic = R. Neoadjuvant XRT; no exposure = False (F), exposure = True (T). Sex; Female = F, Male = M.

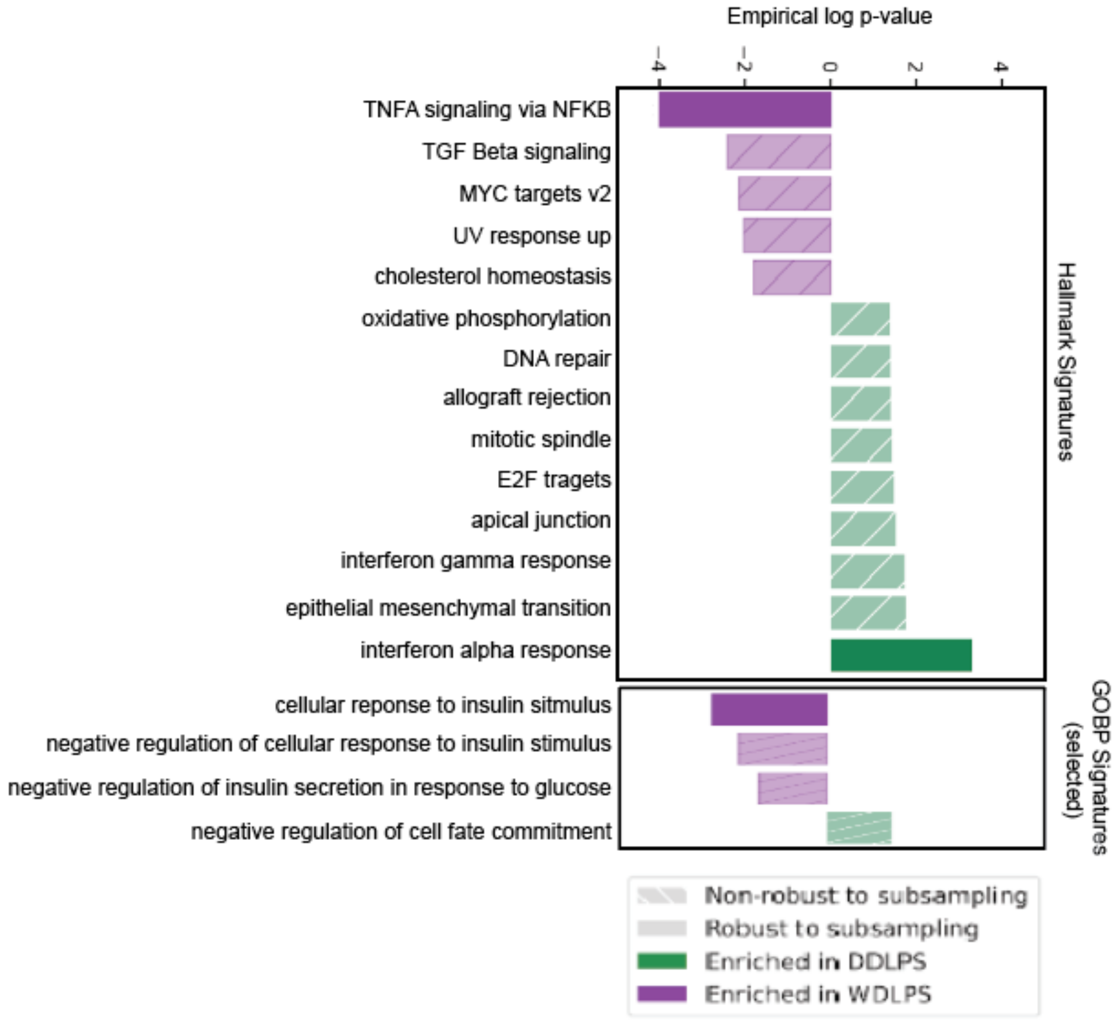


**Figure S6. Robust gene set enrichment analysis.** Output of BEANIE depicted for Hallmark gene sets (top) and selected GO Biological Processes gene sets (bottom). Solid bars indicate pathways enriched in either WDLPS or DDLPS even after accounting for patient specificity and any imbalance in cell numbers between groups.

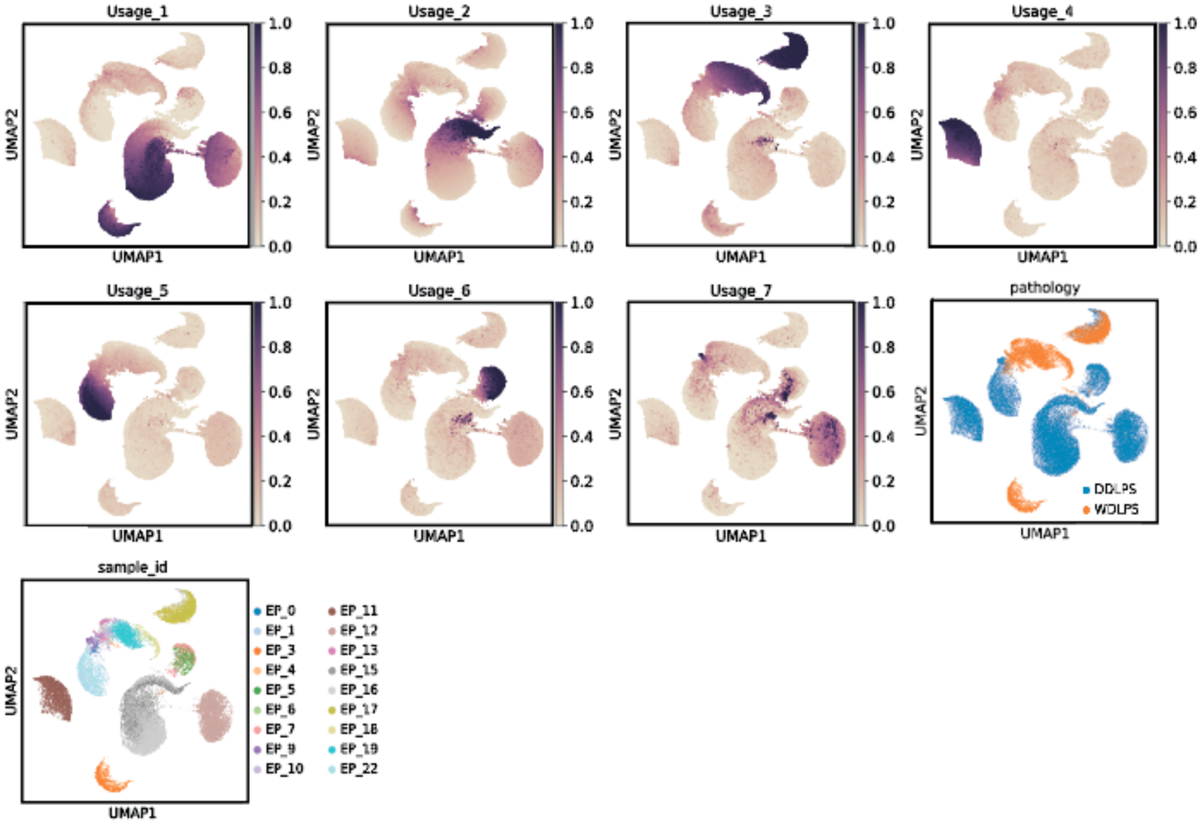


**Figure S7. cNMF usage programs in WDLPS, DDLPS tumor cells.** Seven total usage programs were observed in LPS tumor cells via cNMF analysis. Relative usage (colored by purple) per cell displayed in UMAPs 1-7. Usage_3 and Usage_1 were associated with WDLPS and DDLPS, respectively, as shown in the UMAP of tumor cells colored by pathology (orange, WDLPS and blue, DDLPS). These programs were also present across patient samples (sample_id, cells colored by sample ID).

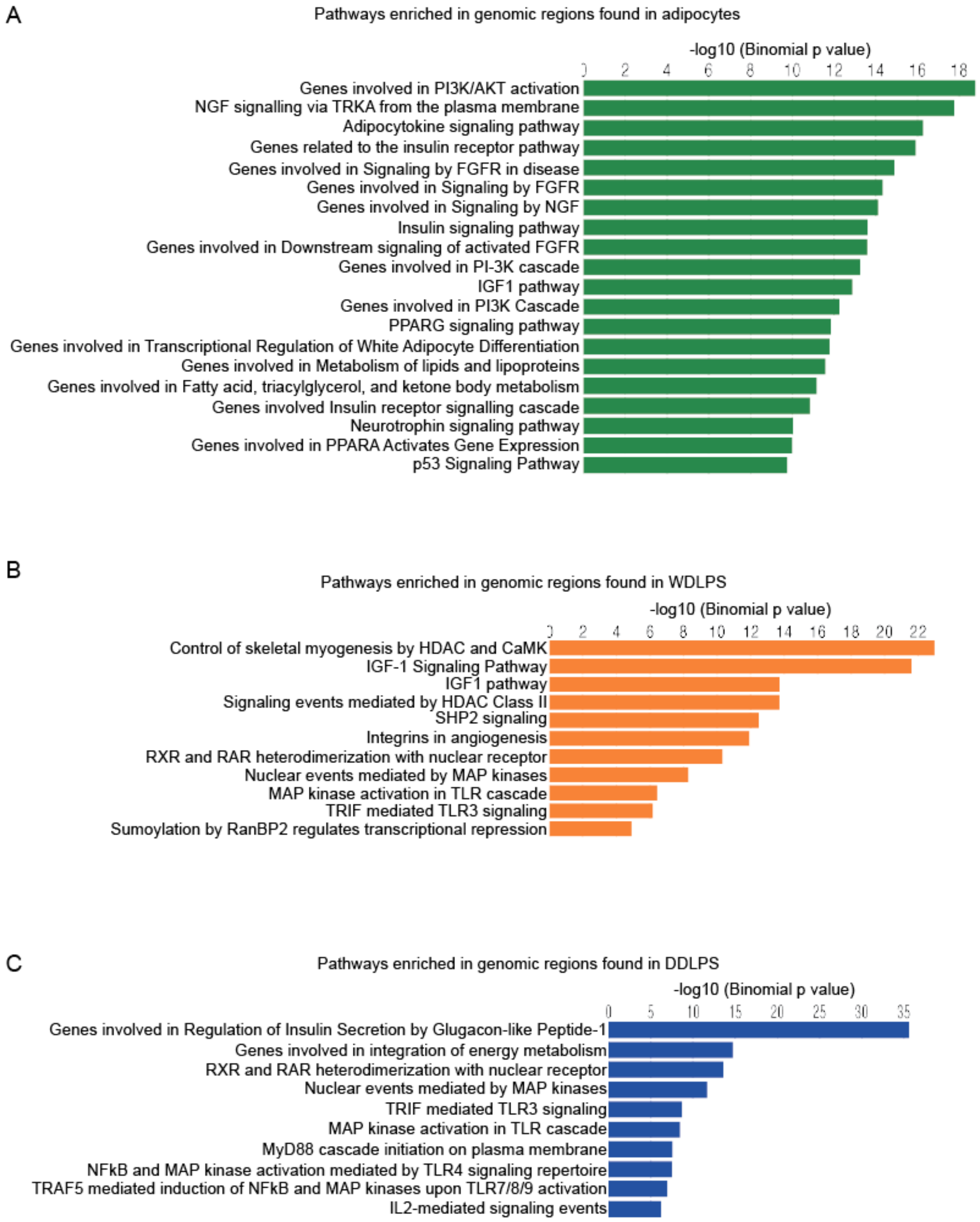

A
Pathways enriched in genomic regions found in adipocytes
-log10 (Binomial p value)
0 2 4 6 8 10 12 14 16 18
Genes involved in PI3K/AKT activation
NGF signalling via TRKA from the plasma membrane
Adipocytokine signaling pathway
Genes related to the insulin receptor pathway
Genes involved in Signaling by FGFR in disease
Genes involved in Signaling by FGFR
Genes involved in Signaling by NGF
Insulin signaling pathway
Genes involved in Downstream signaling of activated FGFR
Genes involved in PI-3K cascade
IGF1 pathway
Genes involved in PI3K Cascade
PPARG signaling pathway
Genes involved in Transcriptional Regulation of White Adipocyte Differentiation
Genes involved in Metabolism of lipids and lipoproteins
Genes involved in Fatty acid, triacylglycerol, and ketone body metabolism
Genes involved Insulin receptor signalling cascade
Neurotrophin signaling pathway
Genes involved in PPARA Activates Gene Expression
p53 Signaling Pathway
B
Pathways enriched in genomic regions found in WDLPS
-log10 (Binomial p value)
0 2 4 6 8 10 12 14 16 18 20 22
Control of skeletal myogenesis by HDAC and CaMK
IGF-1 Signaling Pathway
IGF1 pathway
Signaling events mediated by HDAC Class II
SHP2 signaling
Integrins in angiogenesis
RXR and RAR heterodimerization with nuclear receptor
Nuclear events mediated by MAP kinases
MAP kinase activation in TLR cascade
TRIF mediated TLR3 signaling
Sumoylation by RanBP2 regulates transcriptional repression
C
Pathways enriched in genomic regions found in DDLPS
-log10 (Binomial p value)
0 5 10 15 20 25 30 35
Genes involved in Regulation of Insulin Secretion by Glugacon-like Peptide-1
Genes involved in integration of energy metabolism
RXR and RAR heterodimerization with nuclear receptor
Nuclear events mediated by MAP kinases
TRIF mediated TLR3 signaling
MAP kinase activation in TLR cascade
MyD88 cascade initiation on plasma membrane
NFkB and MAP kinase activation mediated by TLR4 signaling repertoire
TRAF5 mediated induction of NFkB and MAP kinases upon TLR7/8/9 activation
IL2-mediated signaling events

**Figure S8. Genomic region pathway enrichment analysis.** ATAC-seq data was used to calculate differentially accessible peaks (FindAllMarkers) per pathology group, then analyzed with GREAT to identify enrichments in MSigDB pathways. Statistically significant pathway enrichments shown for adipocytes (A), WDLPS (B), and DDLPS (C).

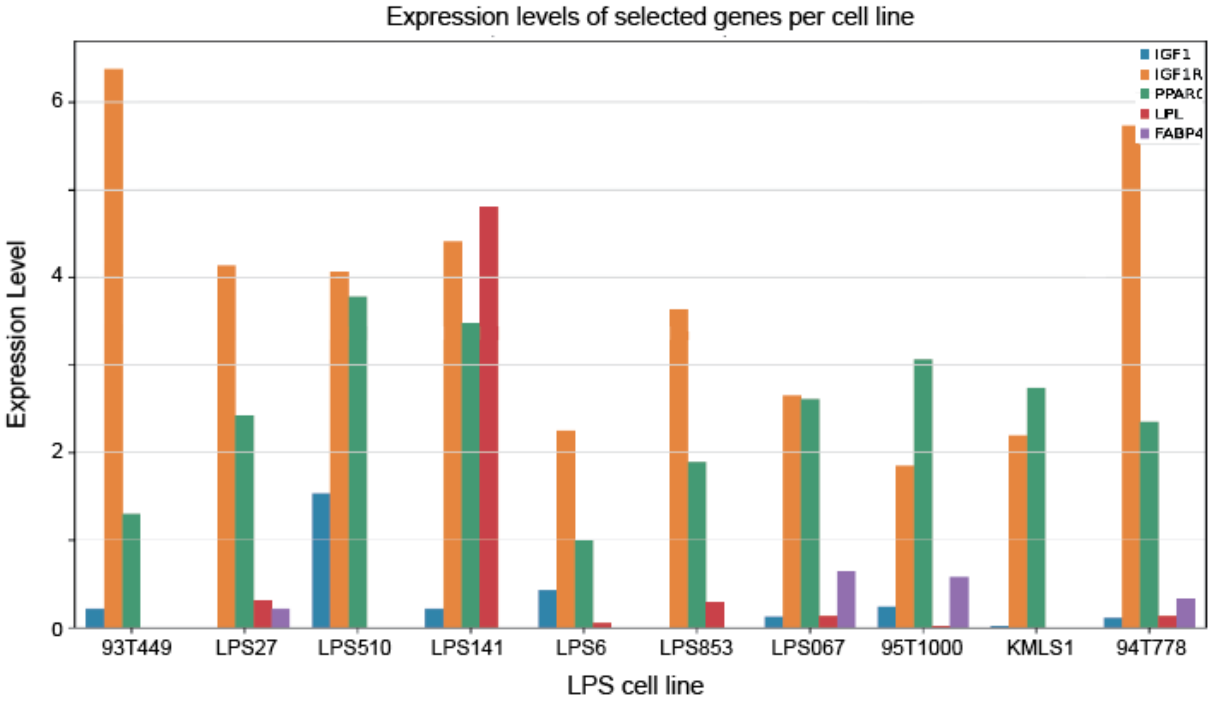


**Figure S9. Expression of IGF1 pathway genes in LPS cell lines.** Expression data derived from bulk RNA-seq data available in the Cancer Cell Line Encyclopedia. Expression values are TPM normalized. The y-axis shows relative expression levels of IGF1 (blue), IGF1R (orange), PPARG (green), LPL (red), and FABP4 (purple) between cell lines, x-axis. The LPS6 and 93T449 cell lines mirrored the pattern of IGF1-associated genes in patient LPS transcriptomic data most closely.

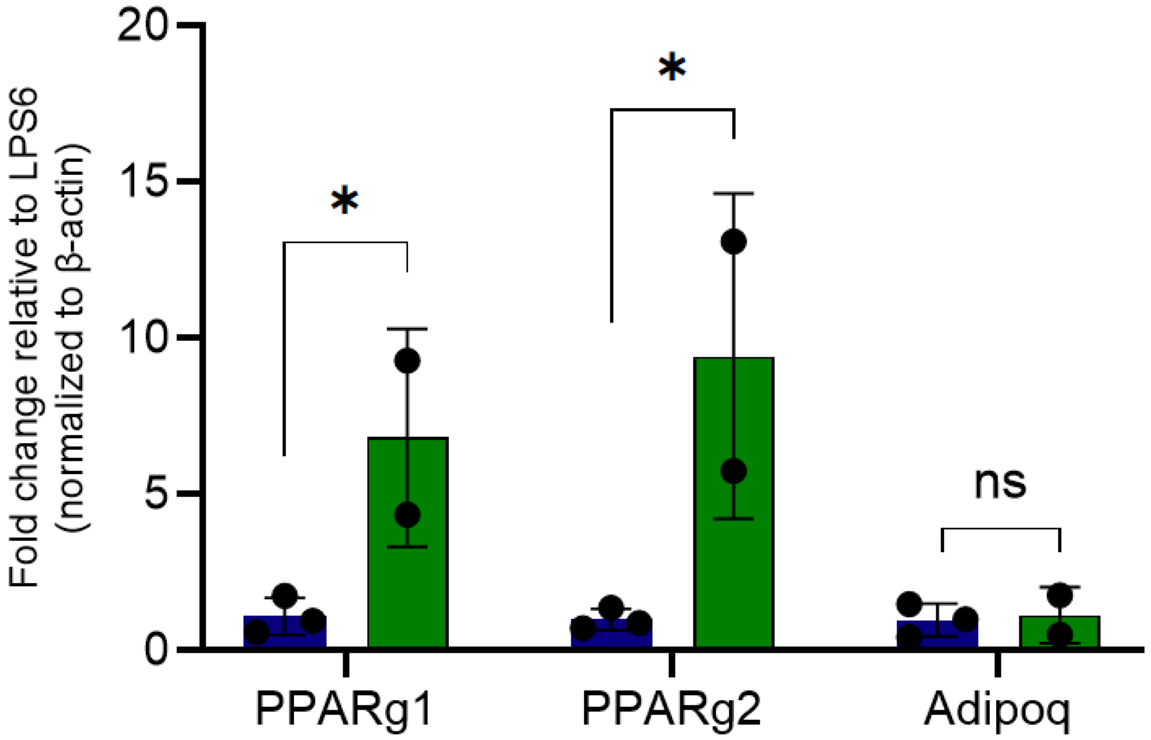


**Figure S10. Relative expression of select PPARG isoforms and ADIPOQ at baseline comparing LPS6 and MSCs.** Fold change relative to expression in LPS6 cells (blue) of PPARG1, PPARG2, and ADIPOQ in adipose-derived human MSCs (green). Expression values are normalized to beta-actin. Statistical significance determined by Student's t-test. *$p < 0.05$, ns = not significant.

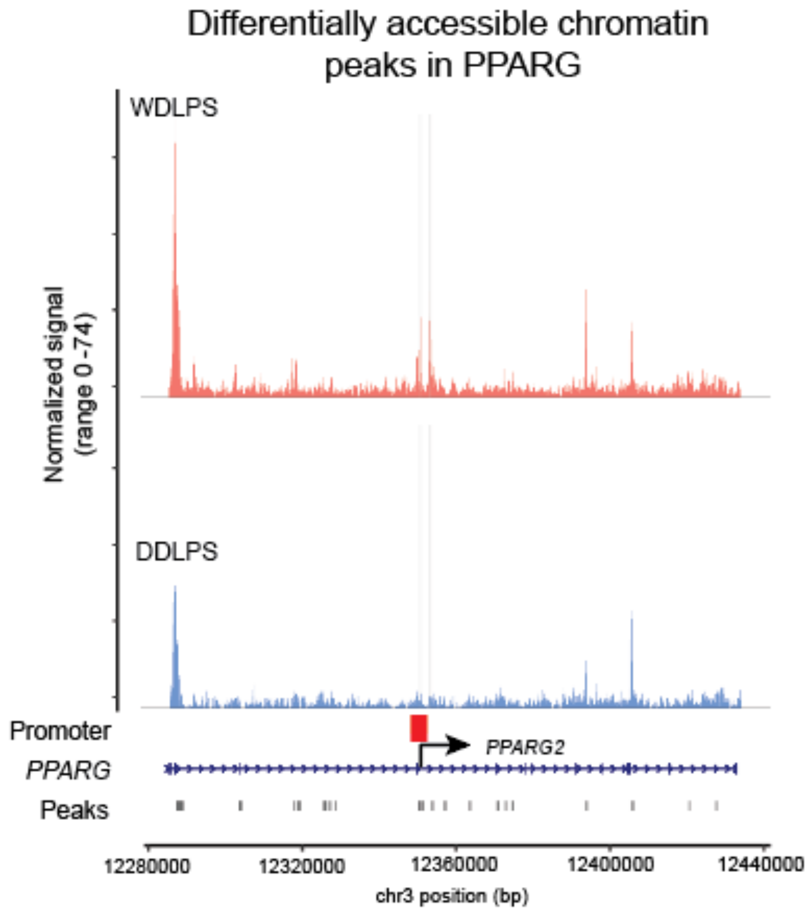


**Figure S11. Differential accessibility over the entire PPARG locus in LPS subtypes.** Coverage plot derived from multiome snATAC-seq data over PPARG gene body, separated by pathology. Peaks highlighted in gray are significantly differentially accessible between WDLPS and DDLPS cells, by logistic regression adjusted for sequencing depth. Significance defined as adjusted p value < 0.05. Promoter-like signature region ('Promoter') denoted by the red bar. PPARG2 TSS denoted by the black arrow. Figure 5C in the main text highlights the region that contains the two differentially accessible peaks within the PPARG gene body.

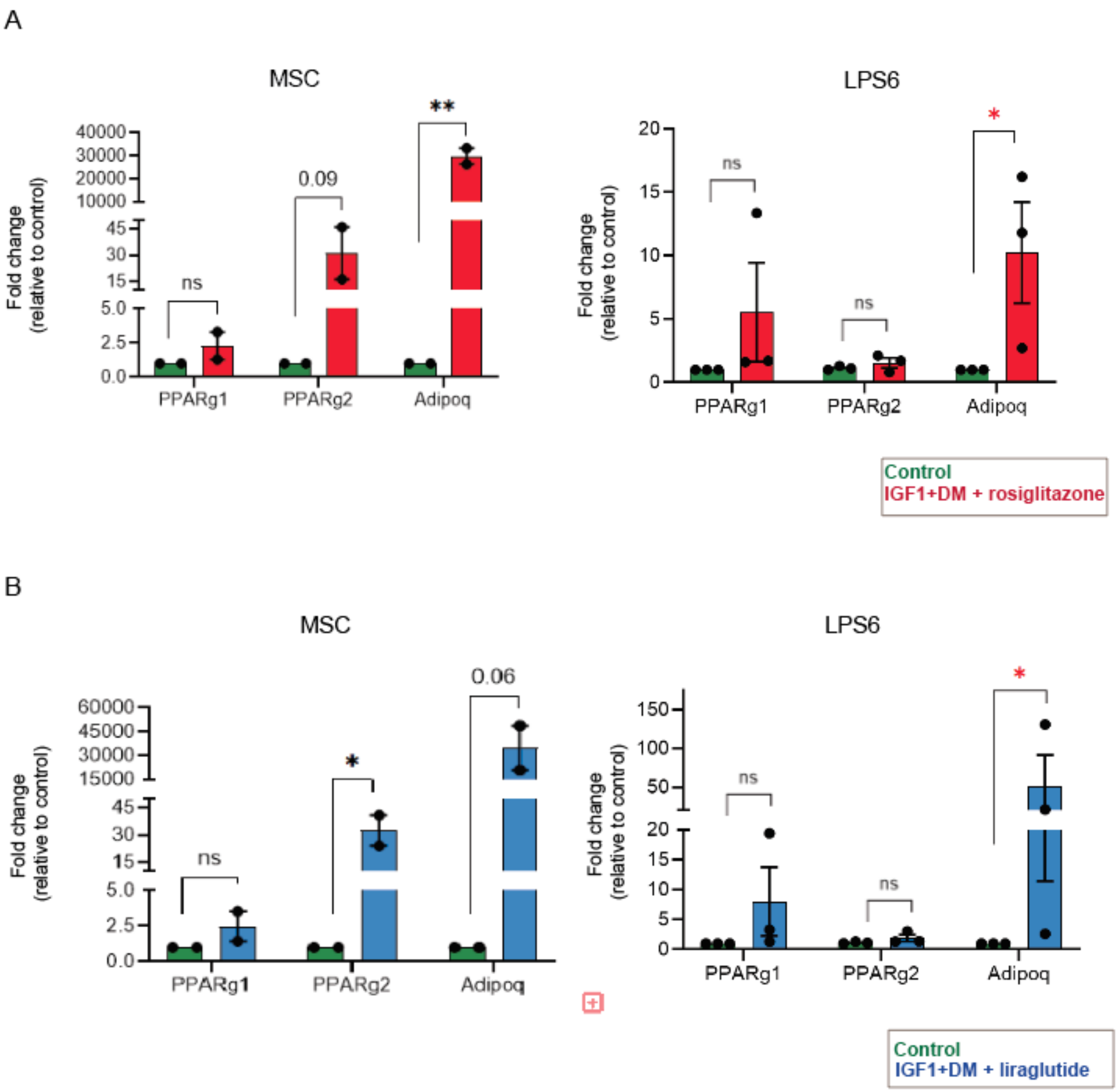


**Figure S12. Differentiation-based therapeutic manipulation of IGF1/PPARG signaling axis.** A. Expression of *PPARG1*, *PPARG2*, and *ADIPOQ* in control cells or cells (green) exposed to IGF1+DM and rosiglitazone (red); MSCs on the left, LPS6 cells on the right. B. Expression of *PPARG1*, *PPARG2*, and *ADIPOQ* in control cells or cells (green) exposed to IGF1+DM and liraglutide (blue); MSCs on the left, LPS6 cells on the right. Statistical significance determined by the Student's t-test. * $p < 0.05$, ** $p < 0.01$, ns = not significant.

**Tables S1 and S2** available in attached Excel file.
